# Cre/*lox*-mediated chromosomal integration of biosynthetic gene clusters for heterologous expression in *Aspergillus nidulans*

**DOI:** 10.1101/2021.08.20.457072

**Authors:** Indra Roux, Yit-Heng Chooi

## Abstract

Building strains of filamentous fungi for stable long-term heterologous expression of large biosynthetic pathways is limited by the low transformation efficiency or genetic stability of current methods. Here, we developed a system for targeted chromosomal integration of large biosynthetic gene clusters in *Aspergillus nidulans* based on site-specific recombinase-mediated cassette exchange. We built *A. nidulans* strains harboring a chromosomal landing pad for Cre/*lox*-mediated recombination and demonstrated efficient targeted integration of a 21 kb DNA fragment in a single step. We further evaluated the integration at two loci by analyzing the expression of a fluorescent reporter and the production of a heterologous polyketide metabolite. We compared chromosomal expression at those landing loci to episomal AMA1-based expression, which also shed light on uncharacterized aspects of episomal expression in filamentous fungi. This is the first demonstration of site-specific recombinase-mediated integration in filamentous fungi, setting the foundations for the further development of this tool.

## Introduction

Filamentous fungi are prolific producers of enzymes and bioactive metabolites with biotechnological applications in pharmaceutical, agricultural and food industries.^1,2^ Importantly, fungal secondary metabolites (SMs) remain a promising source of novel drug leads.^3^ The genes required to produce a SM are usually colocalized in the genome, forming biosynthetic gene clusters (BGCs).^3^ Each BGC contains around 2–20 genes including a large backbone enzyme, such as a polyketide synthase (PKS) or a non-ribosomal peptide synthetase. BGCs can easily be identified in fungal genome sequences and uncharacterized BGCs represent an almost untapped resource for compound discovery.^4^ However, a large fraction of BGCs remain silent or lowly expressed under standard culture conditions, which limits their analysis.^3^ Additionally, as we gain access to the genomic information of diverse fungi promising BGC candidates are often identified in fungal species that are difficult to cultivate or genetically intractable. Hence, heterologous expression of BGCs in hosts with more genetic tools available has become an attractive strategy for genome-based natural product discovery from such fungi.^5^

Filamentous fungi hosts present several advantages for the heterologous expression of BGCs from other filamentous fungi, such as increased compatibility of promoters and intron splicing, and their natural capability for producing SMs.^5–7^ In particular, *Aspergillus nidulans* has been widely used to produce SMs by chromosomal integration or episomal expression of heterologous BGCs.^7^ Our laboratory has been continuing to expand the synthetic biology toolbox for *A. nidulans*, including a tripartite AMA1-based episomal vector system and CRISPRa for silent BGC activation.^8,9^ Episomal systems based on the replicator AMA1 have facilitated the reconstitution of BGCs for SM discovery due to their high transformation efficiency compared to integrative vectors.^10–12^ However, the phenotypic stability of AMA1-vectors has been shown to be limiting.^13^ Therefore, chromosomal expression is preferred for large-scale long-term stable bioproduction in industrial settings.^14^ As strains with chromosomally integrated genes can be grown in low-cost complex substrates, such as agricultural by-products,^15^ as they do not require selection pressures to maintain heterologous genes.

Chromosomal integration of heterologous BGCs can be achieved via random or targeted integration. Given the outcomes of random integration is less predictable and extensive screening is sometimes required to identify producing strains, targeted integration is often preferred.^16^ Currently, targeted integration in *A. nidulans* is pursued through homologous recombination (HR) facilitated by strains deficient in the non-homologous end joining pathway (Δ*nkuA*).^17^ However, HR efficiency drops when integrating constructs larger than a few kilobases.^14^ As a result, HR-mediated integration of large BGCs relies on the laborious sequential integration of smaller BGC fragments.^18,19^ The efficiency of HR can be increased by creating a CRISPR/Cas induced double stranded DNA (dsDNA) break, but multiple integration rounds may still be needed to reconstitute a full cluster.^20–22^ To leverage the increasing amount of BGCs identified in novel fungi, new methods are needed for the effective one-step creation of strains for heterologous BGC expression.^4^

Here, we develop a Cre/*lox* site-specific recombinase system for one-step chromosomal integration of BGCs in the heterologous host *A. nidulans*. Site-specific recombinases are well suited for the integration of large DNA regions, as they mediate the strand exchange between the recombination sites in a size-independent manner and orthogonally to the host machinery.^23^ As Cre/*loxP* recombination is reversible, strategies for irreversible integration rely on 34-bp *lox* sites with mutations either in the 8-bp asymmetric core or 13-bp palindromic ends. Heteromeric *lox* sites contain nucleotide variations in the left or right ends, respectively named LE and RE (Figure 1, Table S1).^24^ Recombination between LE/RE mutant *lox* sites on a chromosomal location and a donor vector will result in a stable integration as the resulting *lox* site on the chromosome will contain both LE and RE mutations and can no longer recombine.^23^ However, on its own LE/RE integration results in the insertion of the whole donor vector (Figure 1). A cleaner strategy for integration relies on two sequential recombination events between two pairs of heterospecific *lox* sites (harboring mutations in the core region) named Recombinase Mediated Cassette Exchange (RMCE). In RMCE the chromosomal landing pad is flanked by two mutually incompatible *lox* sites that can recombine with corresponding compatible sites located in the donor vector flanking the genes of interest (Figure 1). In the presence of the Cre recombinase, the donor vector is first integrated via recombination with one of the *lox* sites at the landing pad. Subsequently, a second recombination event takes place chromosomally between two compatible *lox* sites, resulting in the excision of the donor vector backbone and the original landing pad cassette, leaving behind the genes of interest irreversibly integrated (Figure 1). RMCE has been used for the integration of large constructs in a wide range of cell factories and model organisms.^25–29^

**Figure 1.**
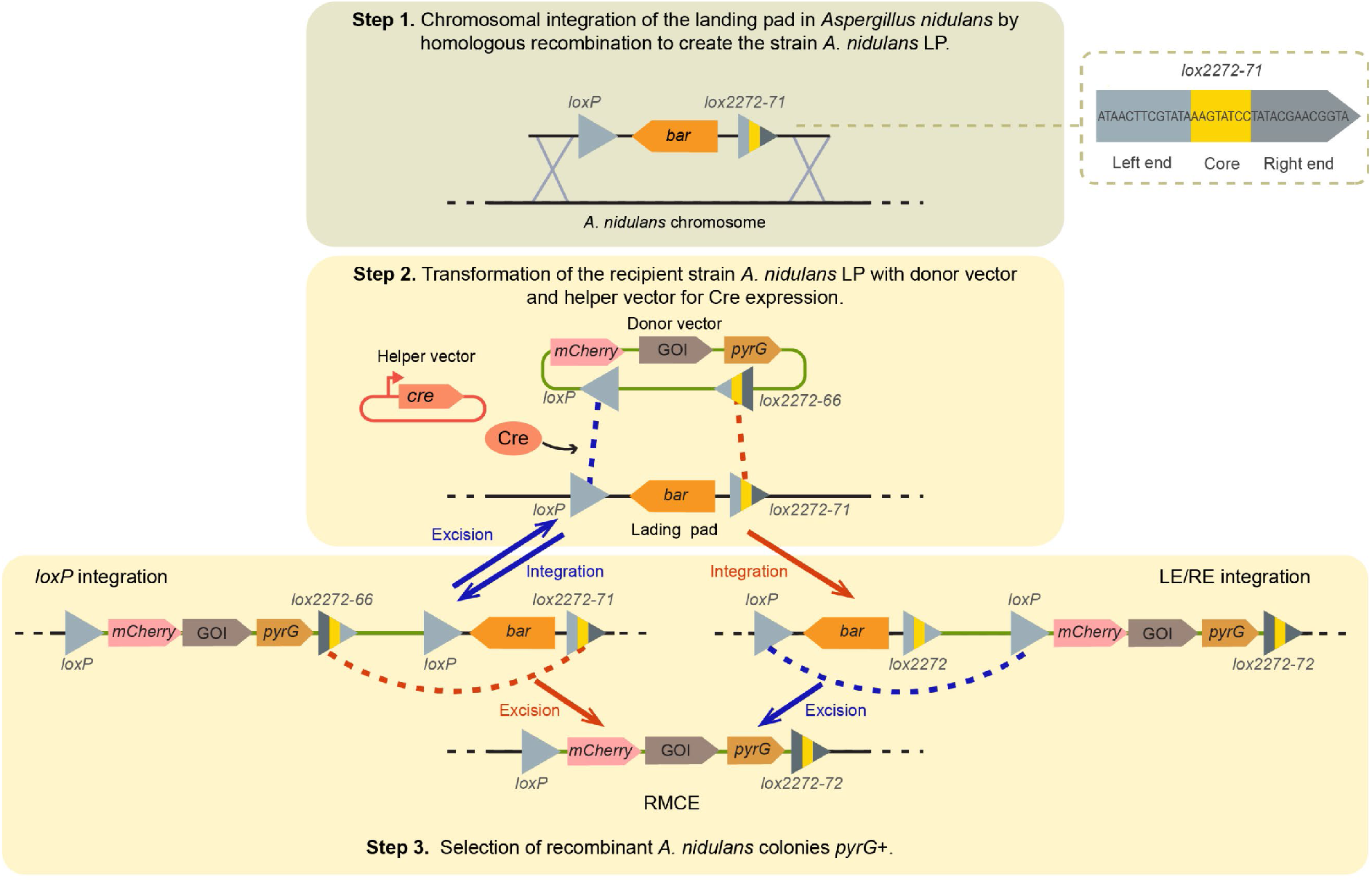
Overview of the strategy for Cre/*lox*-mediated chromosomal integration (not to scale). **Step 1)** The short landing pad (LP) containing the *bar* marker gene flanked by *loxP* and *lox2272-71* is integrated into the destination locus of *Aspergillus nidulans* by homologous recombination, creating the strain *A. nidulans* LP. **Step 2)** *A. nidulans* LP protoplasts are transformed with the donor vector, which contains *loxP* and *lox2272-66* flanking the marker gene *pyrG*, a fluorescent reporter and the genes of interest (GOIs), along with a helper vector for transient expression of Cre recombinase. Cre mediates stable integration by LE/RE recombination or by RMCE in two recombination events (integration and excision). **Step 3)** The recombinant transformant colonies are selected in minimal media for *pyrG* complementation. As the strain *A. nidulans LP* is reused in further transformation rounds, **step 1** is only performed once.

Although Cre/*lox*-mediated deletion has been used to recycle marker genes in filamentous fungi,^30–37^ recombinase-mediated chromosomal integration remained unadopted. Here, we build a vector set for Cre/*lox*-mediated integration in filamentous fungi and demonstrate that this tool is an efficient alternative for the integration of large heterologous BGCs in *A. nidulans*. As a proof-of-concept, we integrate a 21.4 kb DNA fragment containing six genes, including a large polyketide synthase gene, for the heterologous production of preburnettiene B, a recently characterized metabolite from *Aspergillus burnetti*.^38^

## Results

### Design and construction of a recombinase-mediated integration system in *Aspergillus nidulans*

We designed an RMCE strategy involving *lox* sites that contain both the heterospecific mutation of *lox2272* ^39^ and either of the heteromeric (LE/RE) mutations of *lox71* and *lox66*, ^40^ creating the sites *lox2272-71* and *lox2272-66* (Figure 1, Table S1). In principle, *lox2272-71* and *lox2272-66* recombination is irreversible as it results in the double LE/RE mutant site *lox2272-72* (Table S1). Additionally, the sites *lox2272/-72/-66/-71* are incompatible with *loxP* for recombination. Therefore, by flanking the donor cassette and landing pad by *loxP* and *lox2272-66* or *lox2272-71*, RMCE integration can be achieved (Figure 1).

First, we validated the capability of Cre to recombine the sites *lox2272-66 and lox2272-71* by an assay *in vitro* (Figure S1). Next, to create the recipient fungal strain, we chromosomally integrated the floxed (flanked by *lox* sites) landing pad (LP) by homologous recombination in the strain *A. nidulans* LO8030.^41^ The landing pad consisted of the sites *loxP* and *lox2272-71* (Figure 1, Table S1) flanking the marker *bar* for glufosinate resistance. We selected as a first landing locus the sterigmatocystin biosynthetic gene cluster boundaries (Δ*stc*), as we have previously used this locus for the chromosomal expression of heterologous genes.^9^ After glufosinate selection of the transformant colonies and PCR verification, the strain *A. nidulans* landing pad 1 (LP1) was isolated for future tests.

For transient Cre recombinase expression, we created a helper vector unable to replicate in *A. nidulans*. The helper vector encodes a cassette for constitutive expression of Cre under the promoter *gpdA* (*P*_*gpdA*_) and the terminator *trpC* (Figure 1). As the *A. nidulans* LP1 recipient strain carries Δ*nkuA*, which minimizes random integration events, the helper vector presumably would be lost during fungal growth.^17^

To test the feasibility of recombinase-mediated integration at LP1, we built vectors containing donor cassettes with the fluorescent reporter *mCherry* and the *pyrG* marker flanked by *loxP* and *lox2272-66* (Figure 2A, Figure S2A). Donor_vector-1 is a 6.6 kb small vector containing a 3.6 kb floxed donor cassette. Donor_vector-2 is 12.2 kb shuttle vector that supports yeast transformation-associated recombination cloning and contains a 6.6 kb floxed donor cassette that incorporates four cloning sites for the expression of biosynthetic genes under alcohol-inducible promoters^11^ or the cloning BGC genes under native promoters (Figure 2a, Figure S2A). To evaluate the integration of a larger donor cassette, we built donor_vector-2-*bue* containing a 21.4 kb floxed cassette that includes part of the *bue* BGC from *A. burnettii* as proof-of-concept (Figure 2a, Figure S2A). The *bue* BGC has recently been characterized by heterologous episomal expression in *A. nidulans* to produce burnettiene A.^38^ As the complete BGC led to production of multiple metabolite derivatives along with the final product, we decided to evaluate instead the integration of the partial BGC containing *bueA/B/C/D/E/R* (without *bueF*) whose expression results in the production of a single peak corresponding to the penultimate pathway product, preburnettiene B (**1**).^38^ As the *bue* cluster is active in the native host^42^ and episomal expression of *bue* genes under native promoters in *A. nidulans* leads to compound production,^38^ we hypothesized that if the partial *bue* BGC was chromosomally integrated into *A. nidulans* we would observe the production of **1** (Figure S2B).

**Figure 2.**
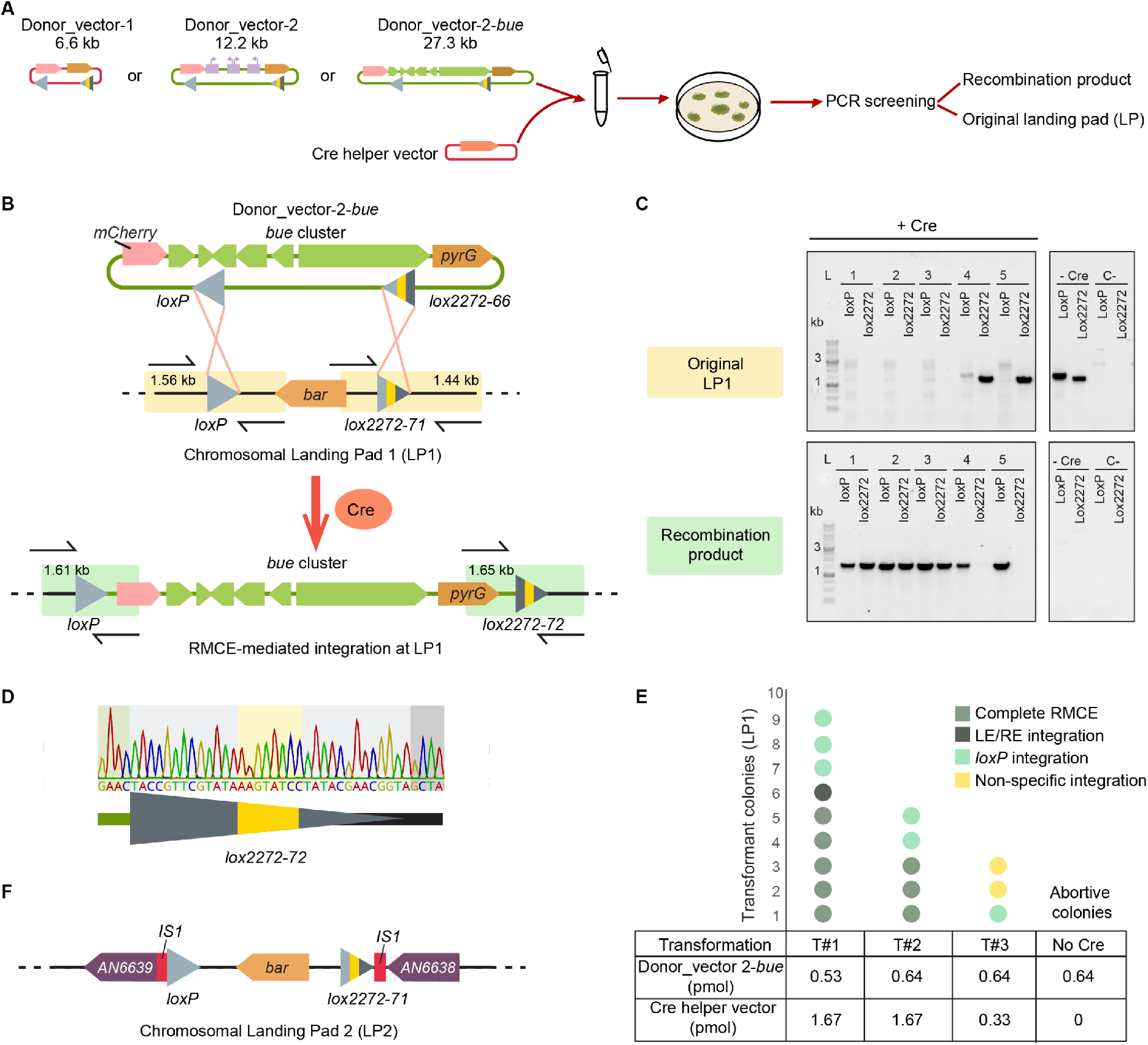
Efficient targeted chromosomal integration of the *bue* biosynthetic gene cluster in *Aspergillus nidulans* **A**. Experimental setup for the evaluation of donor vectors. **B**. Schematic the strategy for *bue* cluster genes integration at landing pad 1 (LP1). PCR amplicons are represented by colored with sizes indicated. **C**. Representative PCR results of transformant colonies for donor_vector-2-*bue* integration. Colonies are indicated as numbers, and the complete gel is found in Figure S3. **D**. Confirmation of the expected recombination event by Sanger sequencing. **E**. Amount of transformant colonies obtained in different transformation experiments with donor_vector-2-*bue*, with the integration mechanism indicated as a color code. **F**. Schematic of landing pad 2 (LP2) in *A. nidulans IS1* locus.

### Cre/*lox*-mediated integration of diverse donor cassettes

To evaluate the efficiency of the recombination system, we transformed protoplasts of *A. nidulans* LP1 by a small-scale PEG mediated transformation in 2 mL microtubes (Figure 2A). We tested different amounts and ratios for each donor vector and the helper vector, and we also transformed *A. nidulans* with the donor vector alone as a control. We consistently obtained viable colonies in the strains transformed with the donor vector along with the helper vector for *cre* expression (3–13 colonies per transformation event) (Table 1). The variation in the number of transformant colonies was independent of the size of the donor fragment, which is expected for recombinase-mediated integration (Table 1).^27^

**Table 1.**
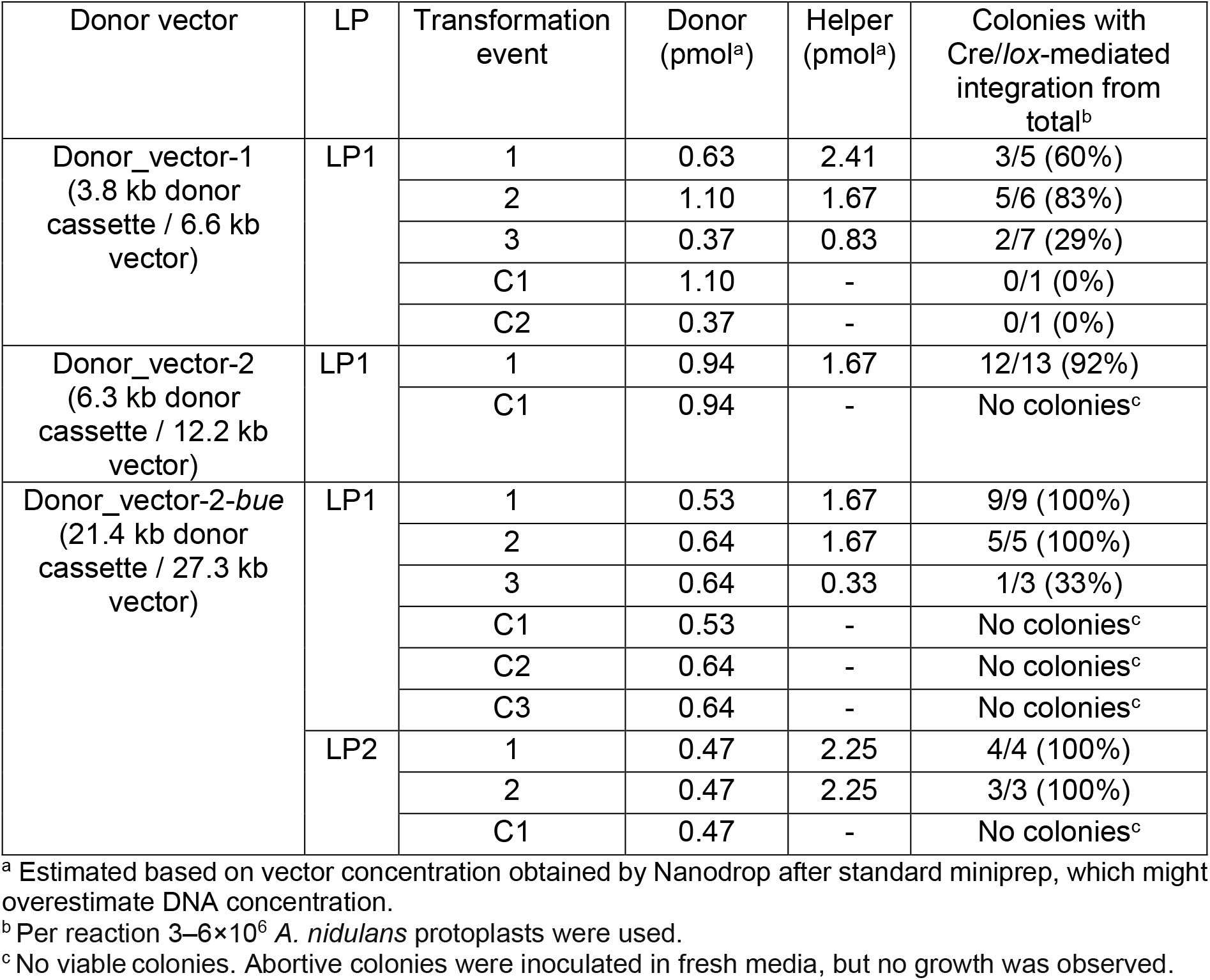
Cre/*lox*-mediated integration experiments in *Aspergillus* nidulans using non-replicative donor vectors, with landing pad (LP) parent strain and negative controls (C) indicated.

In the control strains without *cre* helper vector we mostly observed only small “abortive” colonies that did not support further growth. Abortive colonies arising from residual non-integrated vector encoding *pyrG* have been previously reported in *A. nidulans* Δ*nkuA* strains.^17^ Interestingly, we observed a higher number of abortive colonies when transforming the smaller donor_vector-1 compared to the larger donor_vector-2 and donor_vector-2-*bue*. Additionally, we did not observe viable colonies in the controls when transforming the larger donor vectors without the *cre* helper vector (Table 1).

To evaluate the mechanism of integration in the transformant *pyrG*^+^ colonies, we analyzed the landing locus by PCR amplification of the recombination junction regions with sets of primers for the original landing pad or the recombination product (Figure 2B and C, Figure S3). The frequency of successful recombinase-mediated integration ranged from 29– 100% across experiments (Table 1). We observed that lower recombination efficiency was found in transformations carried out with < 0.9 pmol of helper vector. At higher amounts of helper vector, Cre-mediated recombination efficiency for donor_vector-1 ranged from 60–83% (Table 1). Importantly, the recombination efficiency of the larger vectors donor_vector-2, and 2-*bue* ranged between 90–100% at the highest helper vector concentrations (Table 1, Figures S3D and E). The lower frequency of false positives seems to indicate that larger donor vectors are less prone to random integration. We also observed variability between the type of recombination output (RMCE, LE/RE integration only or *loxP* integration only) with varying donor vector and helper vector amounts (Figure 2E, Figure S3B). At this stage, we also confirmed by sequencing of PCR products that the recombination product between *lox2272-66* and *lox2272-71 in vivo* is the double LE/RE mutant site *lox2272-72* (Figure 2D).

To evaluate the recombination system at a different loci, we created an *A. nidulans* strain harboring a landing pad in the locus *IS1* (LP2), which has been used for heterologous BGC expression (Figure 2, Figure S4A).^18^ When evaluating integration of donor_vector-2-*bue* at LP2 with sufficient *cre* helper vector we obtained 100% efficiency of recombinase-mediated integration in both experiments mostly by RMCE (Table 1, Figure S4B–D).

To investigate if the helper vector is lost in the absence of selection pressure, we analyzed the presence of the *cre* gene by PCR, and obtained positive results in several colonies across experiments (Figure S5). These results could imply that traces of the residual vector might be retained at later growth stages or that random integration of the *cre* helper vector occurs in *A. nidulans*.

### Phenotypic stability of recombinant *Aspergillus nidulans* strains

To evaluate the recombinant strains we first analyzed the expression of a *P*_*gpdA*_*-mCherry* fluorescent reporter encoded in the cassettes integrated at LP1 or LP2. We consistently observed fluorescence in the mycelia of the recombinant strains compared to the negative control (Figure 3A, Figure S6). To benchmark chromosomal expression, we compared the recombinant strains to strains expressing *mCherry* from an episomal AMA1-pyrG vector. We observed that AMA1-based expression was stronger than the chromosomally integrated counterparts when the mycelia was grown under selective conditions, but AMA1-based expression was mostly lost in mycelia grown under non-selective conditions (Figure 3A, Figure S6).

**Figure 3.**
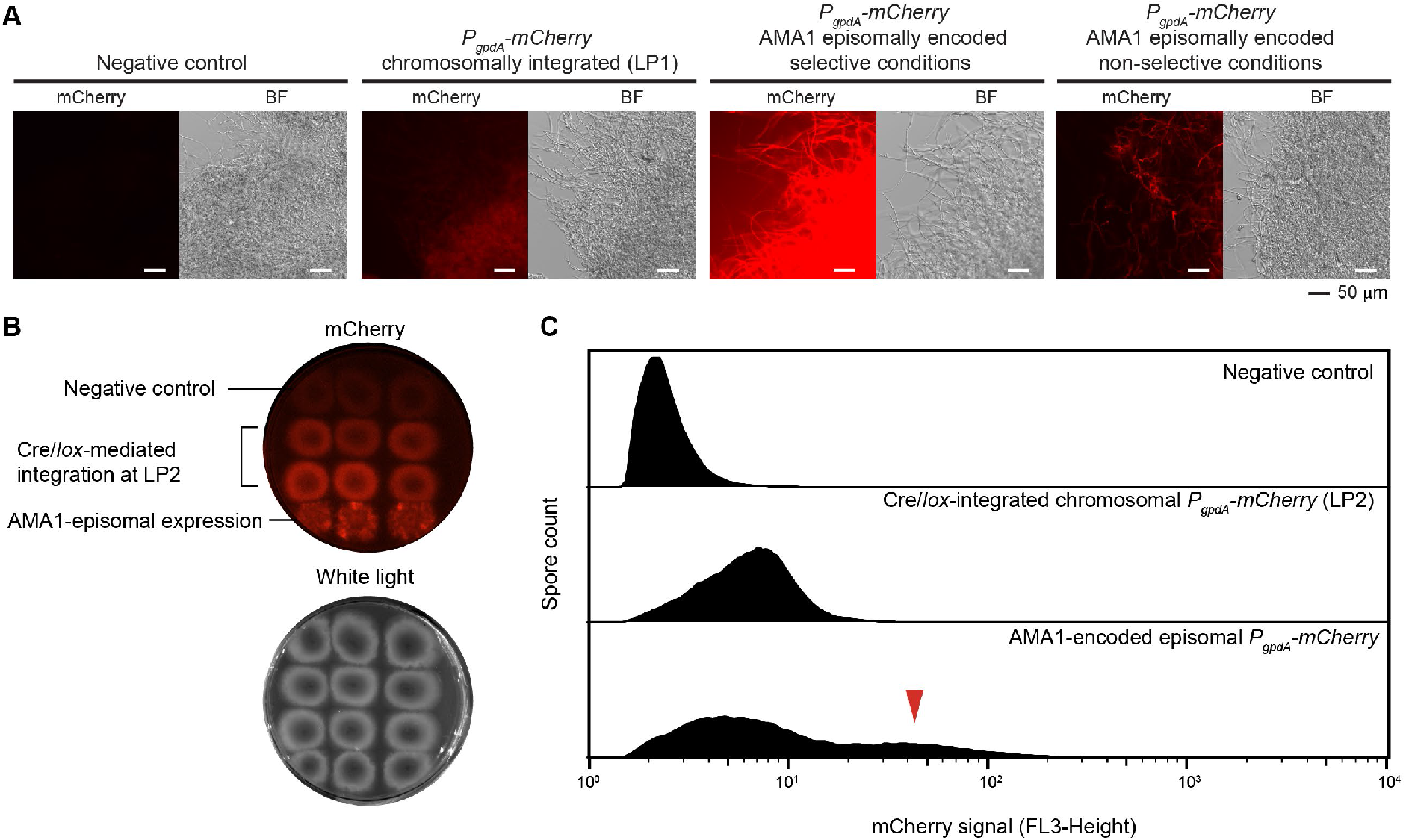
Expression of *mCherry* integrated at the chromosomal landing pads (LPs) compared to AMA1-based episomal expression. **A**. Strains with *mCherry* chromosomally integrated at LP1 show consistent but lower fluorescence than the AMA1-based episomal counterpart under selective conditions. Selection pressure is needed to support high AMA1-based episomal expression. Samples with similar mycelial growth were observed under mCherry filter and brightfield (BF) by fluorescence microscopy. Biological replicates are found in Figure S6. **B**. The expression in recombinant *A. nidulans* colonies at LP2 is more homogeneous than the strains encoding mCherry episomally, which show a patchy expression pattern even under selective conditions. Extended information in Figure S8. **C**. Analysis of spores by flow cytometry shows more compact and homogeneous fluorescence in samples with chromosomal integration of *mCherry* at LP2 compared to AMA1-based episomal expression. A proportion of the spores expressing *mCherry* episomally from AMA1 vectors can reach fluorescence levels one order of magnitude higher than spores with chromosomal expression at LP2 (red arrow). Extended information in Figure S7.

Inspired by comparative studies of chromosomal and episomal expression in yeast by Jensen *et al*,^43^ we analyzed the spores of recombinant colonies by flow cytometry to test phenotypic stability. We observed a unimodal distribution of fluorescence in the strains with *mCherry* integrated at LP1 or LP2, distinguishable from the negative control (Figure 3C, Figure S7). The strains with AMA1-based episomal expression of *mCherry* presented a much wider multimodal distribution, indicative of the heterogenous expression of AMA1-encoded genes on spores even under selective conditions (Figure 3C, Figure S7).^13^ While these results indicated that chromosomal expression at LP1 or LP2 results in a more homogeneous cell population, expression in the recombinant strains was at least one order of magnitude lower than the best performing spores from strains with AMA1-episomal expression (Figure 3C, Figure S7).

To further evaluate the expression pattern of mCherry in spores during fungal growth in solid media, we analyzed the recombinant strains at LP2 by fluorescent photography (Figure 3B, Figure S8). We observed that the recombinant strains for chromosomal expression showed uniform fluorescence in selective and non-selective conditions, distinguishable from the negative control. In contrast, the strains with AMA1-encoded *mCherry* presented a patchy expression pattern across the fungal colony both under selective and non-selective conditions (Figure 3B, Figure S8). Subsequent spore dilutions showed AMA1-based mCherry expression was lost in most colonies under non-selective conditions, but interestingly some colonies in non-selective media retained fluorescence (Figure S8B). Overall, our analysis of the fluorescent reporter at LP1 or LP2 supported phenotypic stability, which is expected of chromosomally integrated genes.

### Expression of the polyketide preburnettiene B from chromosomally integrated genes

To assess the production of the heterologous polyketide **1**, we cultivated recombinant strains with *bueA/B/C/D/E/R* chromosomally integrated at LP1 or LP2. Surprisingly, we did not observe detectable production of **1** when the genes were chromosomally integrated, while we consistently observed the production of **1** in the strains with *bue* genes episomally encoded on AMA1 vector (Figure S9). For troubleshooting, we verified the correct integration of *bue* genes in recombinant strains by whole genome sequencing (Figure S10). Thus, and given that mCherry was still expressed (Figure S9), the lack of compound production could be due to the *bue* genes being silent when integrated chromosomally.

We previously demonstrated that overexpression of the *bue* cluster-specific transcription factor (TF) *bueR* increased the production of burnettienes in an episomal context.^38^ Therefore, we hypothesized that overexpression of the TF *bueR* would activate the expression of the chromosomal *bue* genes in the recombinant *A. nidulans* strains (Figure 4A). Effectively, we observed the production of **1** in strains harboring *bue* genes integrated by RMCE at either LP1 or LP2 when further transformed with an AMA1-based vector encoding *P*_*gpdA*_*-bueR* and the transporter gene *bueG* ^38^ (Figure 4B and C). This result validates the presence of a functional and complete integrated copy of *bueA/B/C/D/E/R*. When comparing strains with *bue* genes integrated at LP1 and LP2, we observed similar production levels between them (Figure 4C).

**Figure 4.**
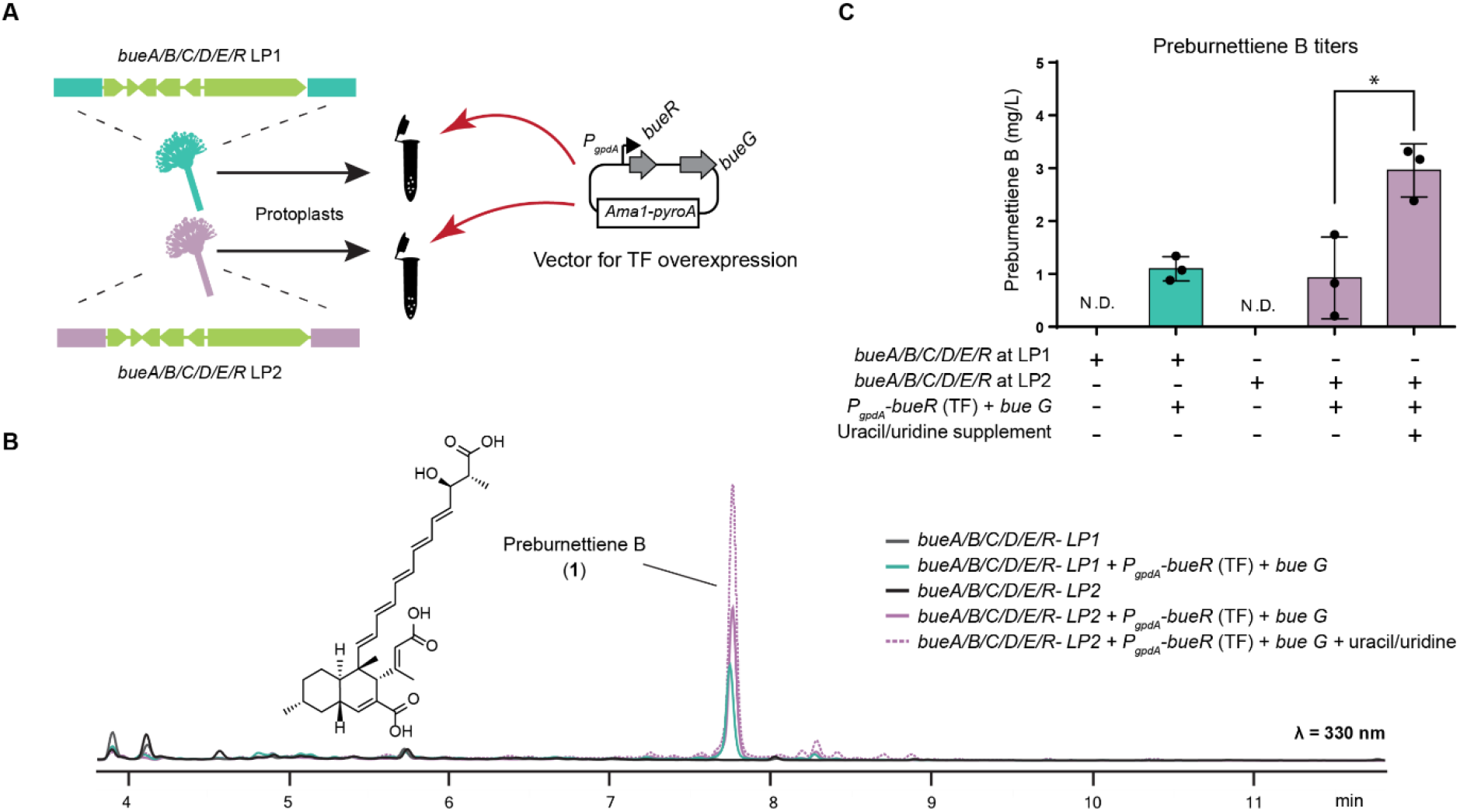
Activation of chromosomally integrated *bue* cluster genes by transcription factor (TF) overexpression. **A**. Protoplasts were prepared from strains with RMCE integrated *bue* genes at LP1 or LP2, and further transformed with a vector for *bueR* overexpression, that additionally encodes the *bueG* transporter. **B**. Chromatogram traces at λ = 330 nm show the activation of the production of preburnettiene B (**1**) in strains with TF overexpression. **C**. Comparable compound titers are observed between the strains with *bue* genes integrated at LP1 or LP2 with TF overexpression on minimal media. Strains grown on media supplemented with uracil and uridine (non-selective condition) presented higher titers than the same strain grown on minimal media (selective condition). Values are the mean of three biological replicates, specific values are indicated as black dots, and error bars represent SD. Two-sided Welch’s T-test p-value = 0.024.

One of the benefits of chromosomal expression is that cultures can be grown under non-selective conditions without compromising productivity. For simplicity, in previous small-scale culturing experiments we cultivated the strains on liquid minimal media with no uracil and uridine, the selective condition of the *pyrG* marker. To evaluate expression under non-selective conditions, we cultivated strains containing the *bue* genes at LP2 and TF overexpression in media supplemented with uracil and uridine. It should be noted that the AMA1-based vector for TF expression is under the selection of pyridoxal auxotrophies with the *pyroA* marker and the medium used lack pyridoxine. As expected, *pyrG* selection pressure was not needed for compound production (Figure 4B and C). Interestingly, we observed a 3-fold higher production in the strains grown in non-selective conditions (Figure 4C). Positional effects arising from insufficient expression of the *pyrG* marker integrated in some loci have been observed in *A. nidulans* before, thus supplementation with uracil and uridine could improve the strain fitness.^44,45^ To confirm that the TF BueR alone was responsible for the activation of *bue* genes, we evaluated strains lacking the transporter *bueG*. Interestingly, we observed even higher production of **1** in the strains with *bueR* overexpression alone compared to the strains additionally expressing *bueG* (Figure S11).

To sum up, overexpression of the cluster-specific TF *bueR* activated the expression of *bue* genes at two different chromosomal contexts. These results demonstrate that recombinase-mediated integration is a feasible strategy to build strains for heterologous compound production, but that alternative strategies might be needed for BGC activation if the gene cluster remains silent. Overall, chromosomal expression permitted cultivating in non-selective media without compromising yields.

## Discussion

In this work we established a system for Cre/*lox*-mediated chromosomal integration of large heterologous BGCs in *A. nidulans*. Site-specific recombinase mediated integration represents a relevant expansion to the synthetic biology toolbox for filamentous fungi, where recombinases had only been used for gene deletion or inversion.^37^ The vector set developed in this work can be adapted for integration at different chromosomal landing loci by replacing the homology arms in the landing pad. Additionally, the vectors are built with promoters and markers portable to other fungi (*P*_*gpdA*_, *T*_*trpC*_, *bar, pyrG*), and therefore could be used to integrate large heterologous DNA in diverse fungal cell factories.^22^

We demonstrated targeted one-step integration of 21 kb DNA regions by RMCE in *A. nidulans*, and this system could be used for integrating larger constructs in future works. We obtained high transformation efficiency (up to 100%) using a small-scale transformation protocol with optimized amounts of helper vector. We additionally observed that the false positive rate and the presence of abortive colonies diminished when transforming large donor vectors (≥12 kb). We also demonstrated that the resulting strains with chromosomally encoded genes presented a more uniform fluorescence phenotype in spores and mycelia compared to AMA1-encoded genes, evidencing genetic and phenotypic stability.

The developed system complements current methods for targeted gene integration in filamentous fungi which rely on DNA repair pathways, such as HR or CRISPR-based knock in (KI). CRISPR-KI has been mostly used to integrate fragments that span few kilobases, and although recently it has been used for the integration of larger BGC fragments in Aspergilli, the efficiency of targeted integration is not reported.^20,21^ In contrast, recombinase-mediated integration does not generate dsDNA breaks and is orthogonal to the host machinery. This makes recombinase-based systems a convenient and reliable alternative for the single-step integration of large DNA constructs, where the efficiency of HR declines. However, recombinase-mediated integration also faces the same constraints as HR for heterologous expression featuring a single gene copy in a chromosomal context. When integrating the heterologous genes *bueA/B/C/D/E/R* under their native promoter at LP1 or LP2 initially we did not observe compound production. Nevertheless, when overexpressing the cluster specific TF *bueR* we restored compound production from the *bue* genes at LP1 and LP2. Strategies for BGC activation, such as TF overexpression, promoter replacement or CRISPRa could be used to activate chromosomally integrated genes that remain silent.^9^ Presumably, integration in a better locus for expression could also result in improved performance.^22^

By benchmarking chromosomal expression to AMA1 based expression, this work also represents an unprecedented characterization of AMA1 episomal expression under the marker *pyrG* in *A. nidulans*. Our original analysis of AMA1-pyrG phenotypic stability by flow cytometry adds a quantitative estimation of the phenotypic heterogeneity of AMA1-based expression in spores from in *A. nidulans* grown under selective conditions in solid media. Even though our results indicate that the phenotypic stability of AMA1-encoded genes in spores is limited, we observed that during mycelial growth in liquid culture there is a more prevalent phenotype for strong expression in both fluorescence and compound production.

Lastly, the Cre/*lox*-mediated integration platform can be expanded in future works. Cre/*lox*-mediated integration could be particularly relevant for integrating large genomic regions containing BGCs cloned by genome capture.^12^ Donor_vector-2 currently supports the assembly of four genes under strong promoters by homology-based cloning, but future donor vectors could be adapted for Type-IIs modular cloning to facilitate the assembly of multigene BGCs where each gene needs promoter replacement.^46,47^ The current system could also be upgraded for simultaneous integration in different loci by using multiple landing pads with different heterospecific recombination sites. To minimize the risk of integration of the helper vector, strategies such as self-excising Cre expression cassettes could also be evaluated.^28^ Additionally, other recombinase systems could be explored for integration in filamentous fungi.^48^

To sum up, Cre/*lox*-mediated integration of BGCs has the potential to speed up the process of constructing strains to produce heterologous metabolites in *A. nidulans*. The capability to uptake and maintain complex exogenous DNA is a key requirement for a good chassis organism for bioproduction.^49^ Thus, this system can be used to upgrade other filamentous fungi chassis. Importantly, this system could be adapted to integrate other large DNA constructs in commonly used industrial Aspergilli for other biotechnological applications.^2^

## Methods

### Vector Construction

All vectors are listed in Table S2 and the oligonucleotides used are listed in Table S4 with their destination vector indicated. Vectors were constructed by isothermal assembly or digestion with restriction site enzymes and ligation. Cloning strategies for each vector are detailed in Supporting Methods. *Cre* was amplified from pBF3038.^50^ Relevant vectors will be made available via Addgene (Table S2).

### *Aspergillus nidulans* strains construction and transformation

Genotype of parental strains are listed in Table S3. *A. nidulans* parental strains with LP1 or LP2 were created by polyethylene glycol (PEG)-calcium-based transformation^51^ with NotI linearized vectors pGemLP1-loxP-bar-Lox2272-71 or pGemLP2-loxP-bar-Lox2272-71 containing 1 kb homology regions for homologous recombination in *A. nidulans* LO8030. Colonies were selected for resistance to glufosinate extracted from Basta as previously^9^, an the event was confirmed by diagnostic PCR.

Protoplasts of *A. nidulans* LO8030 LP1 or LP2 were prepared from germlings,^51^ mixed with a quarter volume PEG 60% to a final concentration of 10^8^ protoplasts per mL and frozen at −80 °C for later use. For *A. nidulans* transformation,, 60 μL of thawed protoplast solution in a 2 mL microcentrifuge tube was incubated with 40 μL of STC buffer (1.2 M sorbitol, 10 mM CaCl_2_, 10 mM Tris–HCl, pH 7.5) and the vector amounts indicated in Table 1. After 20 min of incubation on ice, 400 μL of the calcium PEG 60% mix was added and mixed gently by inversion, followed by a 20 min incubation at room temperature. After adding 1 mL of STC buffer the mix was spread on two plates of stabilized minimal media (SMM) supplemented with pyridoxine (p+) and riboflavin (r+) but lacking uracil or uridine (u-) for auxotrophic selection (u-p+r+), that were then incubated for three days at 37 °C to generate transformant colonies.

Relevant strains of *A. nidulans LO8030-LP1-bueA/B/C/D/E/R* and *A. nidulans LO8030-LP2-bueA/B/C/D/E/R* were analyzed by whole genome sequencing with the DNBSEQ-2000 PE150 platform at BGI Tech Solutions co. (BGI, Hong Kong), resulting in ∼2 Gb of raw genome data (150 bp, paired-end). The reads were mapped to chromosome sequences containing the expected recombination product using Geneious 11.03.

For the activation of the chromosomally integrated *bue* genes by TF overexpression, protoplasts were prepared from the strains *A. nidulans LO8030-LP1-bueA/B/C/D/E/R* and *A. nidulans LO8030-LP2-bueA/B/C/D/E/R*. Protoplasts were transformed with the vector pYFAC-bueG-PgpdA-bueR^38^ or pYFAC-PgpdA-bueR and selected on SMM u-r+p-.

### Diagnostic PCRs

Genomic DNA (gDNA) was extracted from mycelial mass after overnight growth in liquid glucose minimal media (GMM) u-p+r+ at 37 °C. The PCR amplification was conducted by 3 min at 94 °C initial denaturing followed by 27 cycles of 30 s at 94 °C, 30 s at 56.5 °C, 60 s at 72 °C, in a thermocycler with Taq polymerase, using ∼30 ng of *A. nidulans* gDNA in a 10 μl reaction volume. The PCR products were run in a 0.8% agarose gel with 2 μl of the AccuRuler 1 kb DNA RTU Ladder (Maestrogen). All agarose gel images were acquired with a Vilber E-Box VX-2 gel imager with software V15.10. Representative PCRs products were amplified with Pfu polymerase for Sanger sequencing.

### Fluorescence microscopy

Spores from three or more individual colonies were picked and grown overnight at 37° C in small petri dishes containing liquid GMM u-p+r+ or u+p+r+. Fluorescence images were captured at 20x on the epifluorescence inverted microscope Eclipse Ti2 (Nikon), using numerical aperture (NA) 0.75 and Plan Apo λ 20x objective lens (Nikon) and a Camera DS-Qi2 (Nikon) controlled by NIS Elements Advanced Research (Nikon). Fluorescent microscopy was carried out under a mCherry filter set (562/40 nm excitation, 593 nm dichroic beamsplitter, and 641/75 nm emission), using a 400 ms exposure and 1.8x analog gain unless specified otherwise. Images were recorded using NIS-Elements Advanced Research AR V4.60 software package (Nikon).

### Flow cytometry

Spores were collected in water with a sterile loop from colonies grown for 3 days in solid GMM u-p+r+ r. The spore suspension was filtered through a syringe containing sterile cotton to remove residual mycelia and diluted to a concentration of ∼1.10^6^ spores/mL. Data acquisition was performed immediately after spore suspension using a FACSCalibur (BD Biosciences) flow cytometer operated with filtered water as sheath fluid. mCherry signal was observed with a 488 nm excitation laser and the filter FL3 (≥670 nm) and 35,000 events were recorded per sample or events within 4 minutes of run time as collected for the filtered water control. The data was processed using FlowJo V10 software (TreeStar). The output was gated according to forward scatter (FCS) to limit to the size range of spores, an example of the gating strategy is indicated on the Figure S7.

### Fluorescent photography

Plates were analyzed in ChemiDoc MP Imaging System (Bio-Rad) and images recorded with ImageLab software V6.1.0 (Bio-Rad). MCherry images were obtained using the excitation source Green Epi illumination and the emission filter 605/50 nm with exposure time of 0.01 seconds. Bright field was captured with white illumination and a standard emission filter in automatic exposure.

### Metabolic profile analysis by LC-DAD-MS

For each transformant strain, spores from individual colonies were re-streaked individually in a solidified GMM (u-p+r+ or u-p-r+) plate and cultivated for three days at 37 °C. Spores were harvested from plates in 1 mL of 0.1% Tween 80 (Sigma, MO, USA) and approximately 10^8^ spores were inoculated into 250-mL flasks containing 50 mL liquid GMM (u-p+r+, u-p-r+ or u+p-r+) medium. Additionally, ampicillin was added to 50 μg mL^-1^. Cultures were incubated for 4 days with shaking set to 200 rpm and 26 °C. At the end of the culture, 20 mL of media was collected in 50-mL falcon tubes by filtration with Miracloth (Milipore, MA, USA). The metabolites were extracted from the liquid culture with 20 mL of an organic solvent mixture containing ethyl acetate, methanol, and acetic acid (89.5 : 10 : 0.5 ratio). The crude extracts were dried down in vacuo and re-dissolved in 0.3 mL of methanol for LC-DAD-MS analysis.

The analyses of the metabolite profiles were performed on an Agilent 1260 liquid chromatography (LC) system coupled to a diode array detector (DAD) and an Agilent 6130 Quadrupole mass spectrometer (MS) with an electrospray ionization (ESI) source. In all cases 3 μL of the methanol dissolved crude extract was injected. Chromatographic separation was performed at 40 °C using a Kinetex C18 column (2.6 μm, 2.1 mm i.d. x 100 mm; Phenomenex). Chromatographic separation was achieved with a linear gradient of 5–95% acetonitrile-water (containing 0.1% v/v formic acid) in 10 minutes followed by 95% acetonitrile for 3 minutes, with a flow rate of 0.70 mL min^-1^. The MS data were collected in the *m/z* range 100–1000 in negative ion mode and UV observed at DAD λ=330 nm. Peak areas were determined by integration using Masshunter Workstation Qualitative Analysis B.07.00 (Agilent). To quantify preburnettiene B (**1**) samples were compared to a calibration curve prepared with a pure standard (Microbial Screening Technologies) and extrapolated to titers of burnettiene per liter (Figure S12). Statistical analysis was performed with GraphPad Prism 8.3.0 by a Two-sided Welch’s T-test using biological replicates.

## Abbreviations

BGCs: biosynthetic gene clusters
dsDNA: double stranded DNA
CRISPR: clustered regularly interspaced short palindromic repeat
HR: homologous recombination
Kb: kilobase
KI: knock in
LE: left end
LE/RE: Left/Right ends recombination
LP: landing pad
PKS: polyketide synthase
Re: right end
RMCE: Recombinase Mediated Cassette Exchange
SM: secondary metabolite
TF: transcription factor.

## Supporting Information

Supporting Methods: cloning strategy and *in vitro* recombination; Figures S1−12: Biological replicates of fluorescence microscopy and flow cytometry, fluorescent photography, donor vector schematics, sequencing results, LC/MS analysis, PCR gels, *in vitro* recombination, calibration curve; Tables S1−4: *lox* sites used, primers, vectors and strains used in this work.

## Acknowledgments

This project is supported by the Australian Research Council (FT160100233 and DP210102180). I.R. was recipient of an UWA PhD Scholarship. We thank Microbial Screening Technologies (MST) for the strain *Aspergillus burnettii* MST-FP2249 and the preburnettiene B standard. *A. nidulans* LO8030 is a gift from Berl Oakley. We thank Hamideh Rezaee for her help building donor_vector-2, and Julia Grassl (UWA) for helping us to set up the flow cytometer.

## Conflicts of interest

There are no conflicts to declare.

## Author contribution

I.R and Y.H.C conceived the project and wrote the manuscript. I.R. designed and performed the experiments and the data analysis.

## Notes

### Competing Interest Statement

The authors have declared no competing interest.

