## Supporting information for "Cre/*lox*-mediated chromosomal integration of biosynthetic gene clusters for heterologous expression in *Aspergillus nidulans*"

### Supporting Methods

#### 1. Vector Construction

The landing pad vector pGemLP1-loxP-bar-Lox2272-71 was built by a sequential cloning process: pBARGPE1,[1] obtained from the Fungal Genetics Stock Centre, was digested with PmlI and AatII and the site *lox2272-71* was created by ligation of the annealed oligonucleotides AattI-Lox2272-71Rv and AattI-Lox2272-71Fw. A fragment containing the marker *bar* flanked by *lox2272-71* and a *loxP* from pBARGPE1 was PCR amplified and cloned by isothermal assembly with NEBuilder HiFi DNA Assembly Master Mix (New England Biolabs, MA, USA) into NotI digested pGemT (Promega) incorporating at the 5′ and 3′ 1 kb homology arms for LP1 amplified from *Aspergillus nidulans* gDNA, creating the vector pGem-LP1. To build pGemLP2-loxP-bar-Lox2272-71, the floxed *bar* cassette was amplified from pGemLP1-loxP-bar-Lox2272-71, and two fragments containing the homology arms for LP2 were amplified from *A. nidulans* gDNA and cloned with NEBuilder HiFi DNA Assembly Master Mix into NotI digested pGemT.

*Cre* was amplified from pBF3038 [2] and cloned into pBARGPE1-LIC.[3] The $P_{gpdA}$-*Cre*-$T_{trpC}$ cassette was then further amplified and cloned into pGemT creating pGem-PgpdA-Cre-TtrpC. The donor vector was initially built using a NotI and MscI digested pKW20088,[4] amplifying a ~200 bp fragment containing *loxP* from pXP322 [2] and creating the *lox2272-66* with a gBlock, resulting in the intermediate vector pKW20088-loxP-pyrG-lox2272-66. On parallel, the coding sequence of *mCherry* was amplified from pMP7601 [5] and cloned into pBARGPE1 under $P_{gpdA}$. pKW20088-loxP-mCherry-pyrG-lox2272-66 was created after PacI digestion of pKW20088-loxP-pyrG-lox2272-66 and isothermal assembly of a PCR amplicon containing $P_{gpdA}$-*mCherry*-$T_{trpC}$. pGem-loxp-lox71-2272 was created amplifying the floxed *pyrG* cassette from pKW20088-loxP-pyrG-lox2272-66 with primers that added NotI sites in the ends of the amplicon and cloning by NotI digestion and ligation to NotI digested pGemT (Promega). pGem-loxp-mcherry-lox71-2272 (donor_vector-1) was created with PacI digestion of pGem-loxp-lox71-2272 and amplification of $P_{gpdA}$-*mCherry*. pRecomb-loxp-mcherry-4Gcloningsite-lox71-2272-66 (donor_vector-2) was created *de novo* by isothermal assembly of PCR parts containing the floxed cassette of pKW20088-loxP-pyrG-lox2272-66, the 4-gene cloning cassette from pYFAC-CH2 (Addgene ID 168978) [6] and parts from pKW20088[4] to include the sequences for plasmid maintenance and selection in *E. coli* and *Saccharomyces cerevisiae*.

pRecomb-loxp-mcherry-BueABCDER-lox71-2272-66 (donor_vector-2-*bue*) was created by digestion Donor2 with NotI and isothermal assembly of four PCR fragments containing the partial *bue* cluster, amplified from *Aspergillus burnettii* MST FP2249 gDNA.

#### 2. *In vitro* validation of *lox* sites

The substrate DNA fragments were PCR amplified with primers LoxP-pyrG-F and GBlock-R from template vector pKW20088-loxP-pyrG-lox2272-66 and with primers LIC-H1-F and PbarUpstream-R-Seq from pGemLP1-loxP-bar-Lox2272-71. The *in vitro* recombination was performed following the protocol for Cre (NEB, M0298M), with 1.30 h incubation at 37 °C and heat inactivation. As control, the PCR fragments were subject to the same treatment but without recombinase. The reaction products were run in a 0.8% agarose gel with 2 µl of the AccuRuler 1 kb DNA RTU Ladder (Maestrogen) and the image was taken with a blue LED transilluminator and a phone camera.

#### Supporting Figures

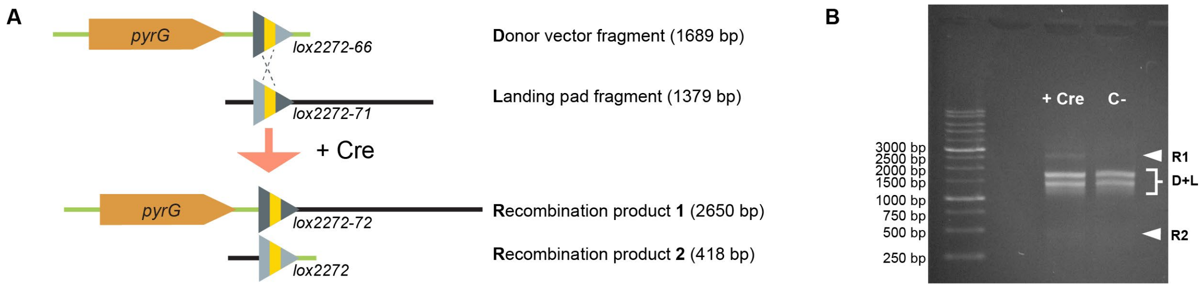


**Figure S1.** *In vitro* validation of the LE/RE mutant sites *lox2272-66* and *lox2272-71*. **A.** Experimental set up. Fragments were amplified by PCR containing the site *lox2272-66* (D) or *lox2272-71* (L). After *in vitro* recombination mediated by Cre, two recombination products are formed of different length compared to the substrates of recombination. **B.** Gel electrophoresis run after *in vitro* recombination reveals a new band of the size of recombination product 1 (R1) in the sample treated with Cre recombinase compared to the control. Recombination product 2 (R2) is less evident due to its smaller size and mass. According to the instructions from the manufacturer (NEB), the efficiency of *in vitro* Cre recombination is low for which this assay is a qualitative approximation. Nevertheless, the larger recombination product was observed validating the designed *lox2272-66/71* pair.

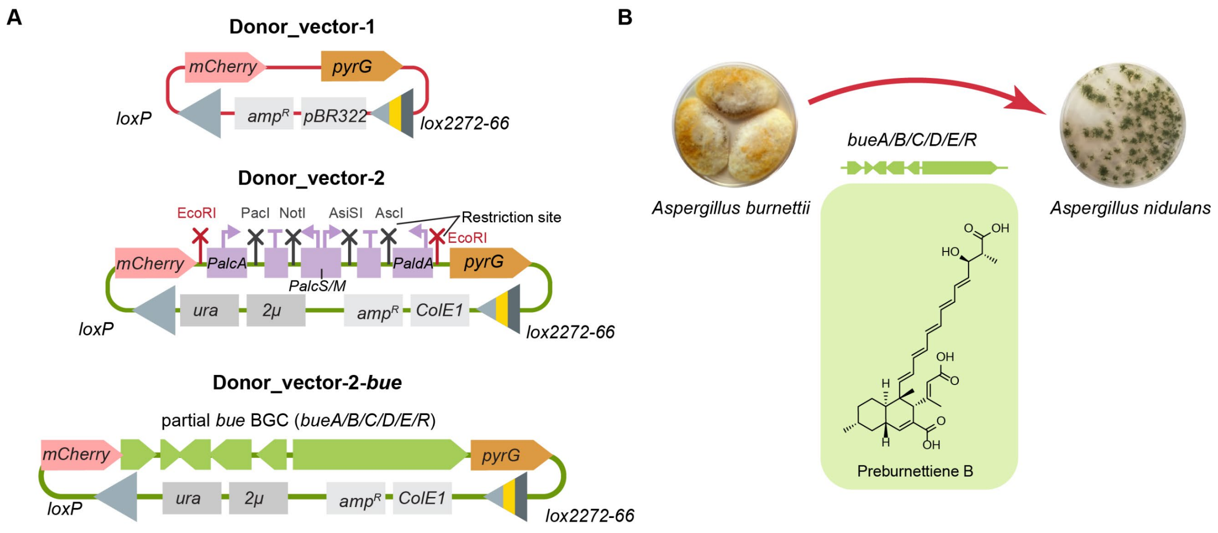


**Figure S2.** Donor vectors evaluated for Cre/*lox*-mediated integration.Donor_vector-1 is a small 6.6 kb vector used in initial testing. Donor_vector-2 is a 12.3 kb shuttle vector with components for replication and selection on *Saccharomyces cerevisiae* (*ura*, *2μ*) and *Escherichia coli* (*ampR*, *ColE1*). Donor_vector-2 contains four alcohol inducible promoters for heterologous expression derived from the vector pYFAC-CH2[6] as well as EcoRI sites for cloning large gene fragments under native promoters (red restriction sites). As an example, we cloned an 18 kb region of *Aspergillus burnetti* containing genes from the *bue* BGC, resulting in a 27.3 kb vector. **B.** Overview of proof-of-concept integration of a fragment from the *bue* biosynthetic gene cluster from *Aspergillus burnetti* in *A nidulans*, which encodes the pathway intermediate decalin polyketide preburnettiene B.

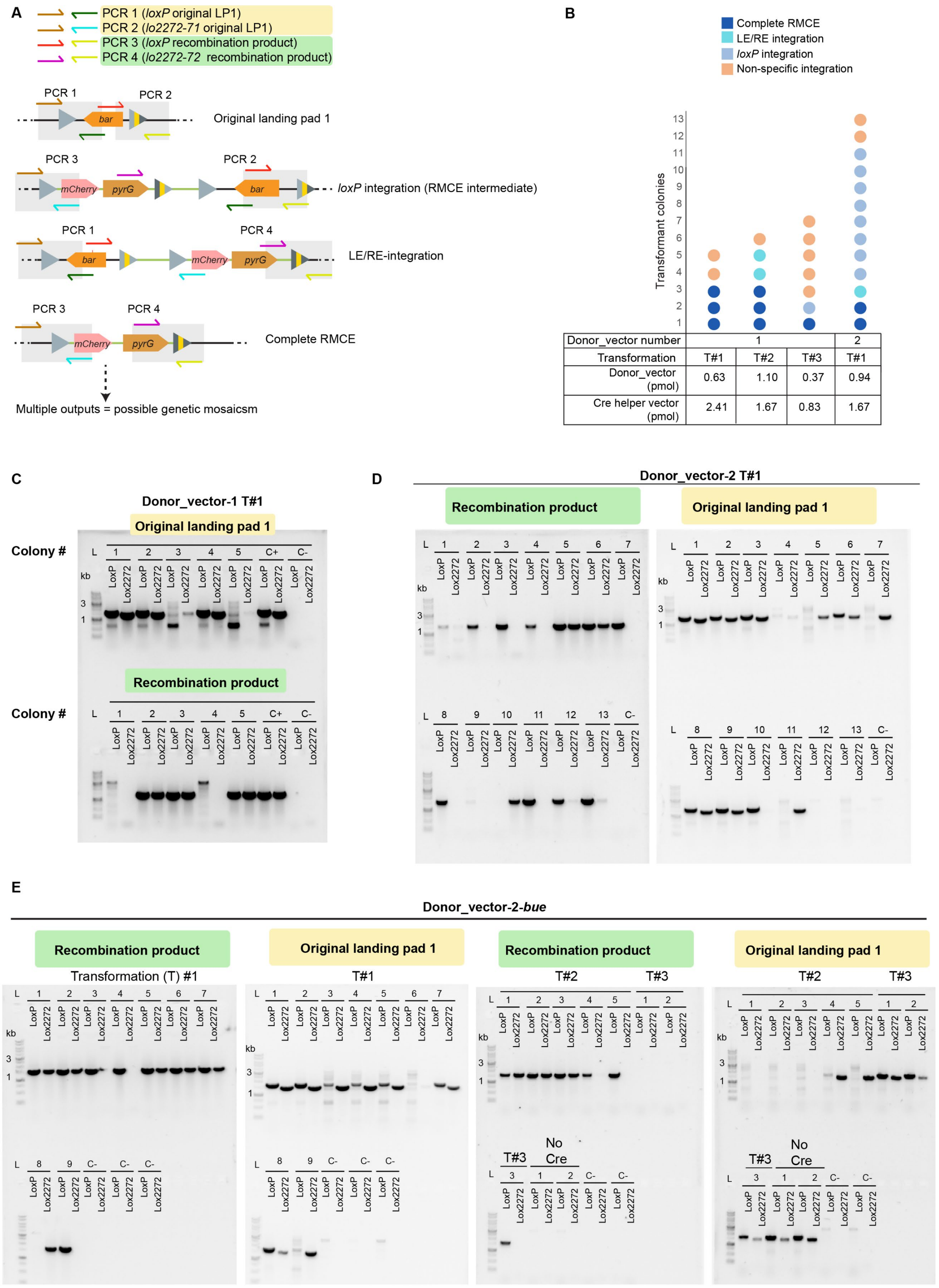


**Figure S3.** PCR-based screening of recombinant colonies. **A.** Overview of the PCR screening strategy. The gDNA from each transformant colony is analyzed by four PCRs to determine the possible outputs. **B.** Amount of transformant colonies obtained in different transformation experiments with donor_vector-1 and donor_vector-2, with the integration mechanism indicated as a color code, as analyzed by PCR. Recombination is obtained across a different amount and proportion of *cre* helper vector and donor vector. **C.** Representative gel of the PCR results of a transformation event with donor_vector-1, different transformant colonies are indicated as numbers. **D.** Representative gel of the PCR results of a transformation event with donor_vector-2, different transformant colonies are indicated as numbers. **E.** PCR results of a transformation events with donor_vector-2-*bue*, different transformant colonies are indicated as numbers. **Note:** Interestingly, a portion of the transformants yielded positive PCR results for the recombination product as well as the original landing pad. Thus, our results could be indicative of genetic heterogeneity in some transformants. If a recombination event happened after the first nuclear division, it could result in heterokaryotic transformants, that can be subsequently isolated by repeating the selection pressure selection.

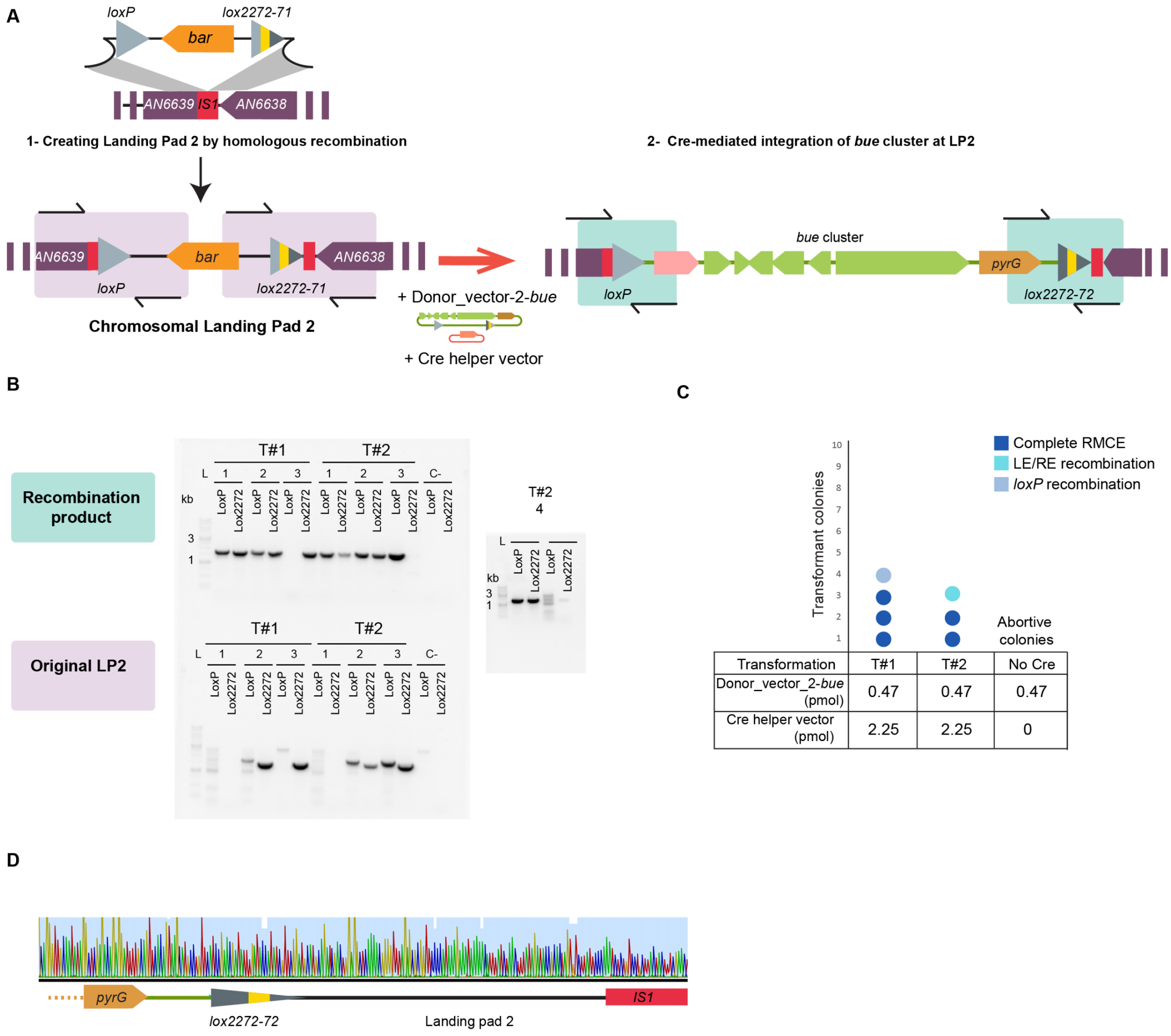


**Figure S4.** Evaluation of integration of donor_vector-2-*bue* at landing pad 2 (LP2). **A.** Schematic overview of the experimental setup to build and test LP2**. B.** PCR screening results of two different transformation events with donor_vector-2-bue, different transformant colonies are indicated as numbers. **C.** Number of transformant colonies obtained in different transformation experiments at LP2, with the integration mechanism indicated as a color code, as analyzed by PCR. **D.** Sanger sequencing of representative PCR amplicon, confirming the site *lox2272-72* after recombination at LP2.

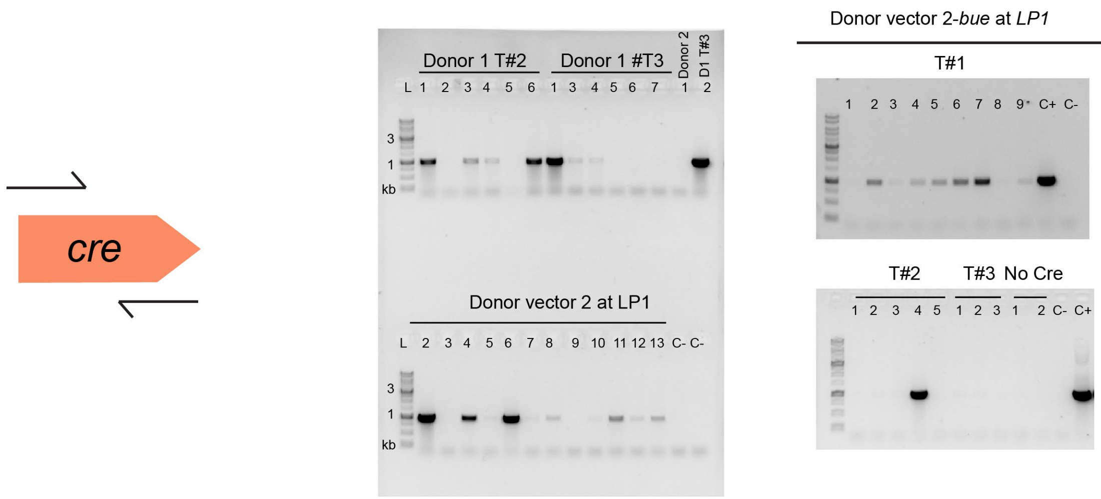


**Figure S5.** Screening for presence of *cre* expression cassette. PCR positive results are found in several transformant colonies with high variability across experiments (0% – 60% colonies per transformation event).These results seem could imply that traces of the residual vector might be present at later growth stages or that that random integration of the *cre* helper vector might occur in some strains.

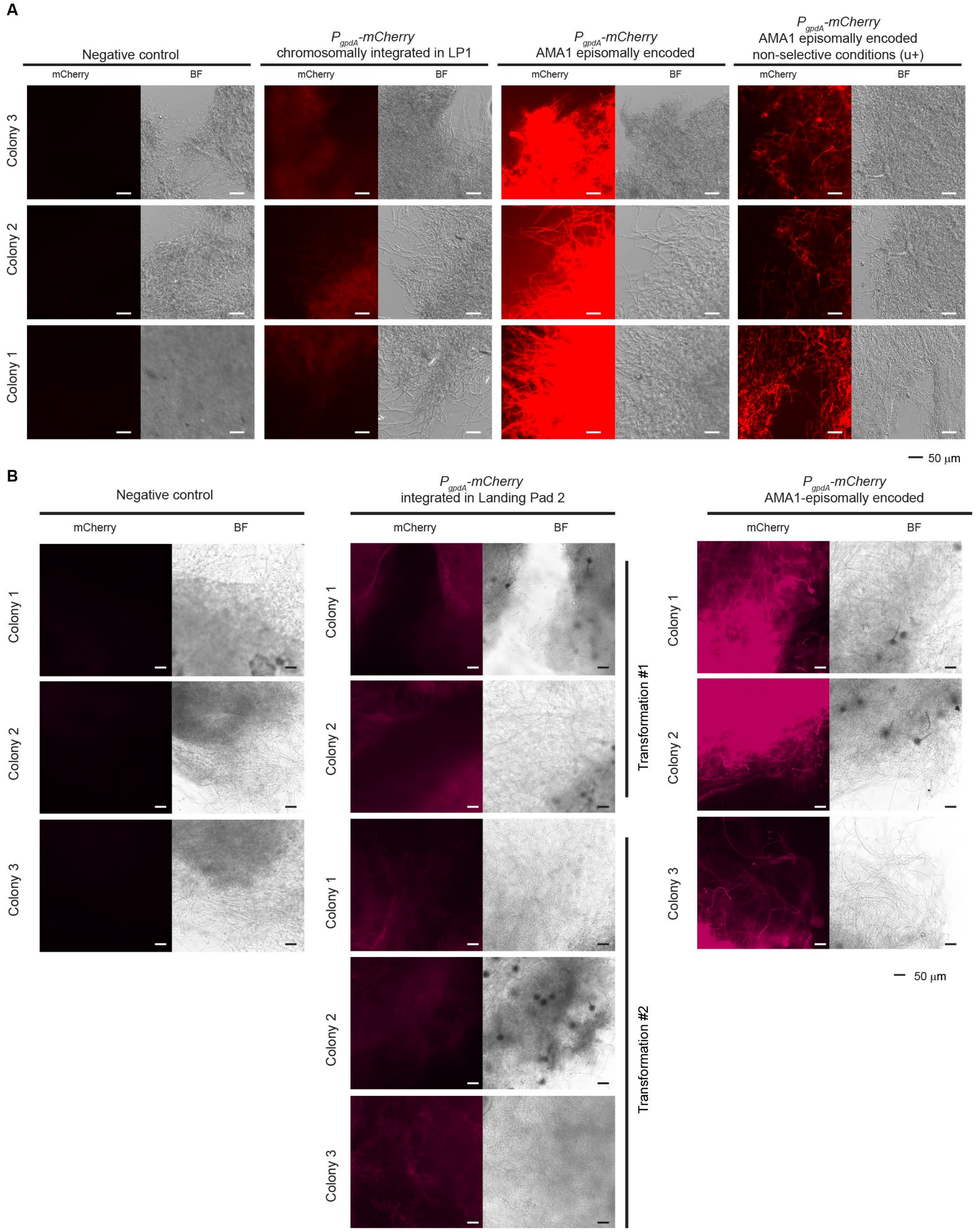


**Figure S6.** *P*$_{gpdA}$*-mCherry* expression in recombinant strains at LP1 or LP2, compared to episomal expression. **A.** Strains with *P*$_{gpdA}$*-mCherry* integrated at LP1 show low *mCherry* expression in mycelia, but distinguishable from the negative control. Expression at LP1 is lower than episomal expression when growing on selective media (lacking uracil and uridine). AMA1-based episomal expression in mycelia grown under non-selective conditions (supplemented uracil and uridine) shows a sparce pattern, but some hypha still show higher signal than strains with chromosomal expression at LP1. The spores for each sample were collected from three individual colonies representing biological replicates of a recombinant strain and grown overnight in liquid stationary culture at 37 °C. **B.** Integration of the donor_vector-2-*bue* at LP2 results in consistent but low mCherry expression in mycelia, but distinguishable from the negative control. Expression at LP2 is lower than episomal expression under selective conditions. All samples were grown on selective media unless specified otherwise. The spores from each sample were collected from individual transformant colonies, and different transformation events when indicated, and grown overnight in liquid stationary culture at 37 °C. Samples with similar mycelial growth were observed under mCherry filter and brightfield (BF). Scale bar 50 μm.

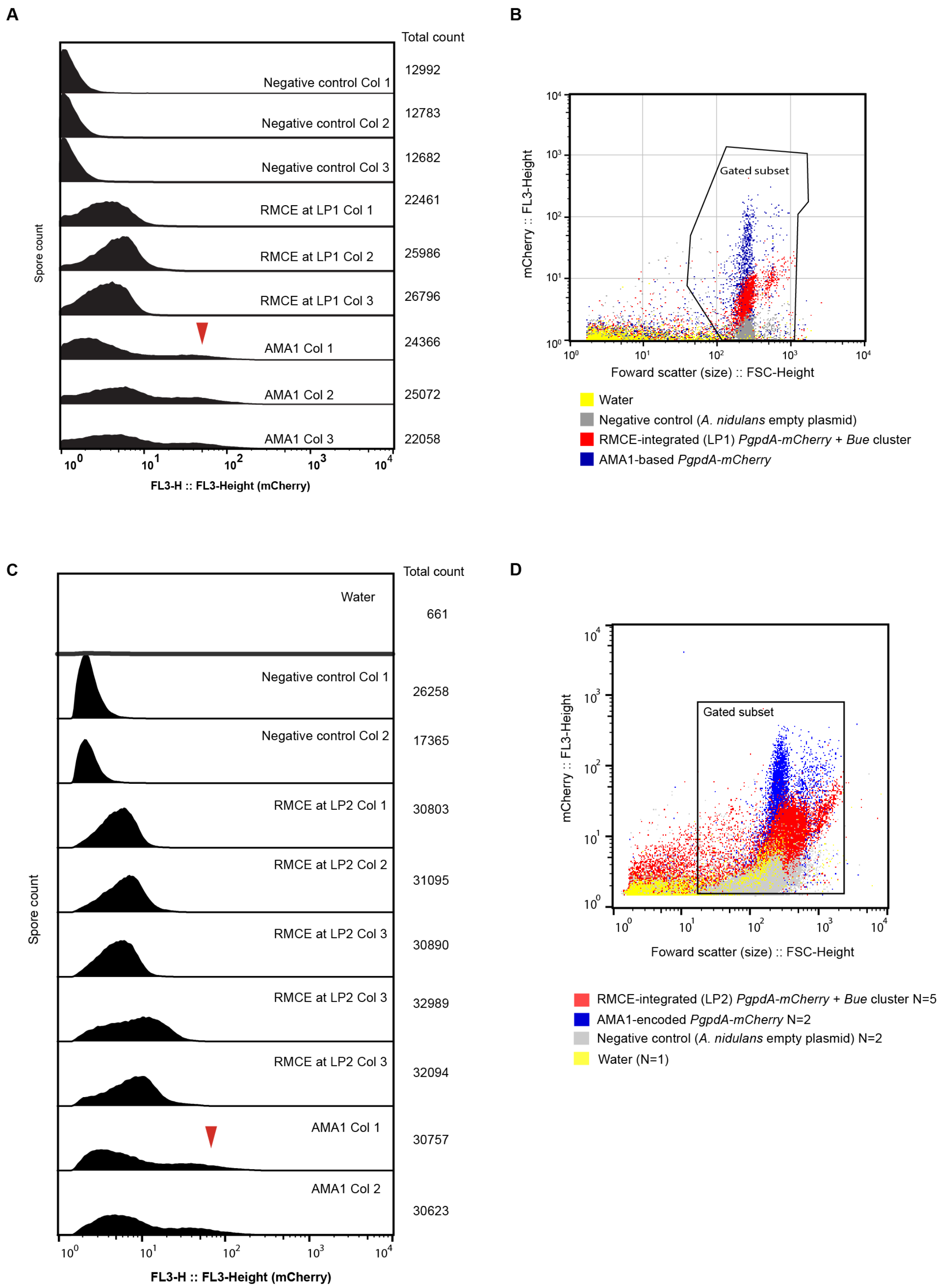


**Figure S7.** Analysis of mCherry in spores by flow cytometry with biological replicates. **A.** Analysis of expression at LP1. Histograms show spore count vs FL3-Height which is indicative of mCherry signal. We observe that the samples with $P_{gpdA}$-*mCherry* integrated at LP1 show fluorescence distinguishable to the negative control, and a more compact distribution than AMA1-based expression. However, a proportion of the spores from colonies with AMA1-based mCherry expression can reach fluorescence levels at least one order of magnitude higher than spores with chromosomal expression at LP1 (see red arrow). Gated event count numbers are indicated at the right. **B.** Ungated plots showing the total of events and gating strategy for analysis at LP1. *y* axis represent mCherry signal and x axis represents particle size. **C.** Analysis of expression at LP2. Histograms show spore count vs FL3-Height which is indicative of mCherry signal. We observe higher signal in the strains with chromosomal integrated mCherry than the control *A. nidulans* strain (empty plasmid). However, signal at LP2 (IS1) is also lower than the higher range of signal from AMA1-based mCherry expression (red arrow). **D.** Ungated plots showing the total events for the analysis of LP2. Gating strategy is indicated.

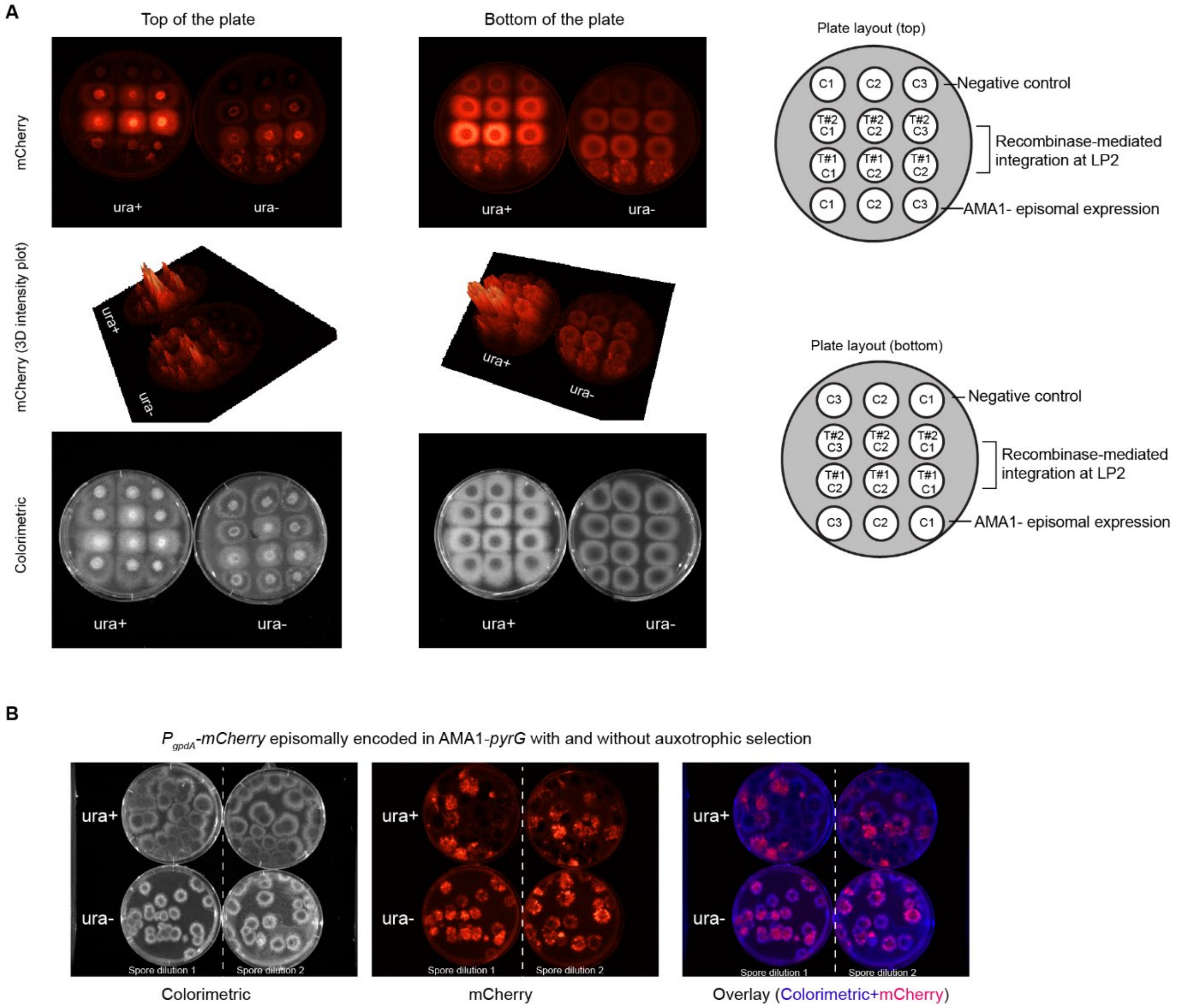


**Figure S8.** Visual inspection of fluorescent phenotype of integrative and episomal $P_{gpdA}$-*mCherry* in *A. nidulans* colonies. **A.** Colonies growing under non-selective (ura+) and selective (ura-) conditions. Images were acquired from both the bottom and the top of the plate. Plate layouts are shown on the right indicating the transformation event and the colony number. Images taken with green light for excitation an mCherry filter show uniform fluorescence in the strains with $P_{gpdA}$-*mCherry* chromosomally integrated, compared to the controls. The strains with episomal AMA1-encoded $P_{gpdA}$-*mCherry* show a patchy pattern of fluorescence. The mCherry signal 3D intensity plot is included to aid the interpretation of mCherry signal. The colorimetric images (white light) are on the bottom. Plates were incubated at 37 °C for three days. **B.** Difference in genetic stability and fluorescence pattern observed in *A. nidulans* colonies product of plating a spore dilution containing AMA1-pyrG encoded $P_{gpdA}$-*mCherry*. As each colony arises from an individual spore, we can observe that under selective conditions all colonies retain fluorescence. Under non-selective conditions, many of the colonies do not show fluorescence. Interestingly, many of the fluorescent colonies under non-selective conditions demonstrate fluorescence comparable to the colonies under selection condition. We also observe more colonies in the plates with no selection pressure (~30 colonies) than in the plates with selection pressure (21-25 colonies) , which is indicative of the genetic stability of AMA1-pyrG. Plates were incubated at 37 °C for three days.

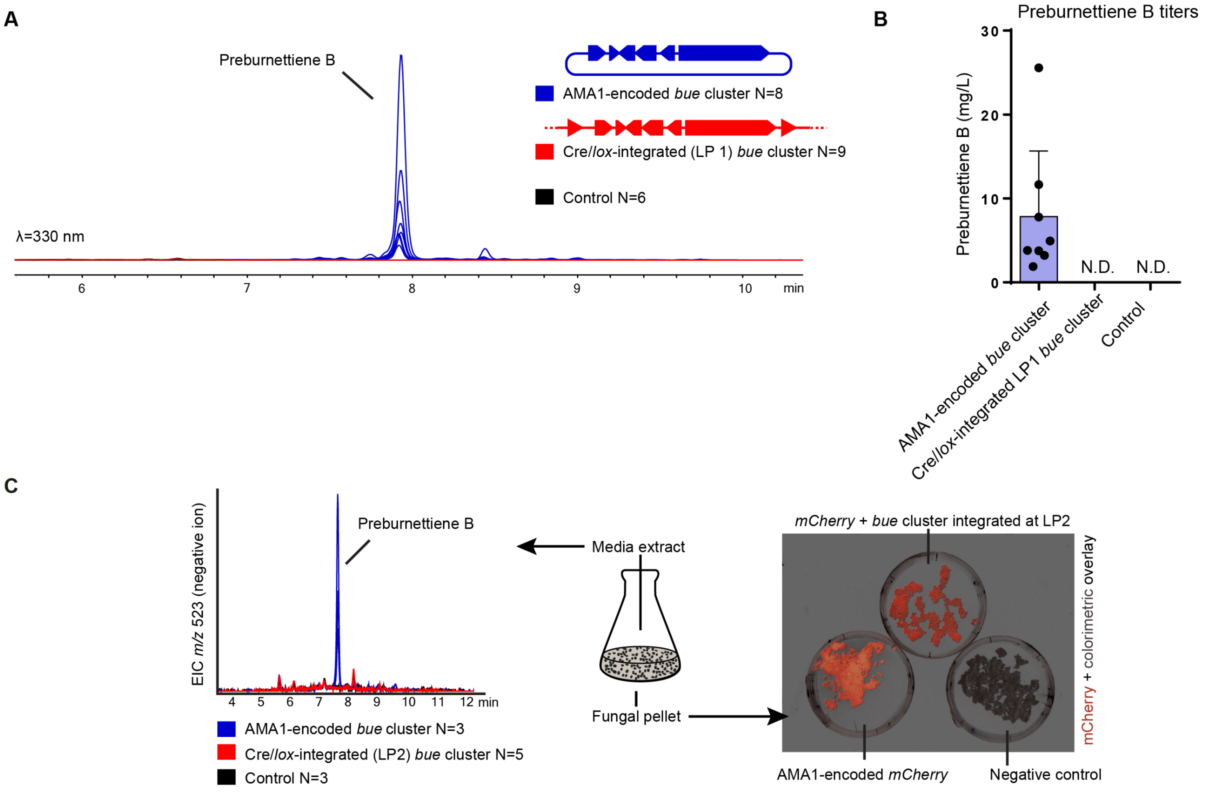


**Figure S9.** Evaluation of preburnettiene B (**1**) production in recombinant strains. **A.** DAD (λ=330 nm) chromatograms of culture media extracts of *A. nidulans* encoding the genes *bueA/B/C/D/E/R* on AMA1 episomal vectors show production of **1**, unlike the strains where *bueA/B/C/D/E/R* are chromosomally integrated at LP1 and in the negative control. **B.** Integrated peak area of the chromatogram in A, showing the variability in compound titers between biological replicates harboring AMA1-based vectors, and lack of production in the strains with the *bue* genes chromosomally integrated. Values are the mean of biological replicates, represented as dots. Error bars are standard deviation. N.D.= Non detected. **C.** (Left) Extracted ion chromatogram on negative ion mode for *m/z* 523 ± 0.5 shows the expected peak for **1** in the strains with AMA1-encoded bue cluster genes (blue) but not in the strains with chromosomally integrated bue cluster genes (red) or the control (black). (Right) Fungal pellets from the end of the culturing period were filtered an observed by fluorescence photography. Fluorescence was observed in the strains with AMA1-encoded $P_{gpdA}$-*mCherry* and strains with $P_{gpdA}$-*mCherry* chromosomally integrated at LP2.

**A**

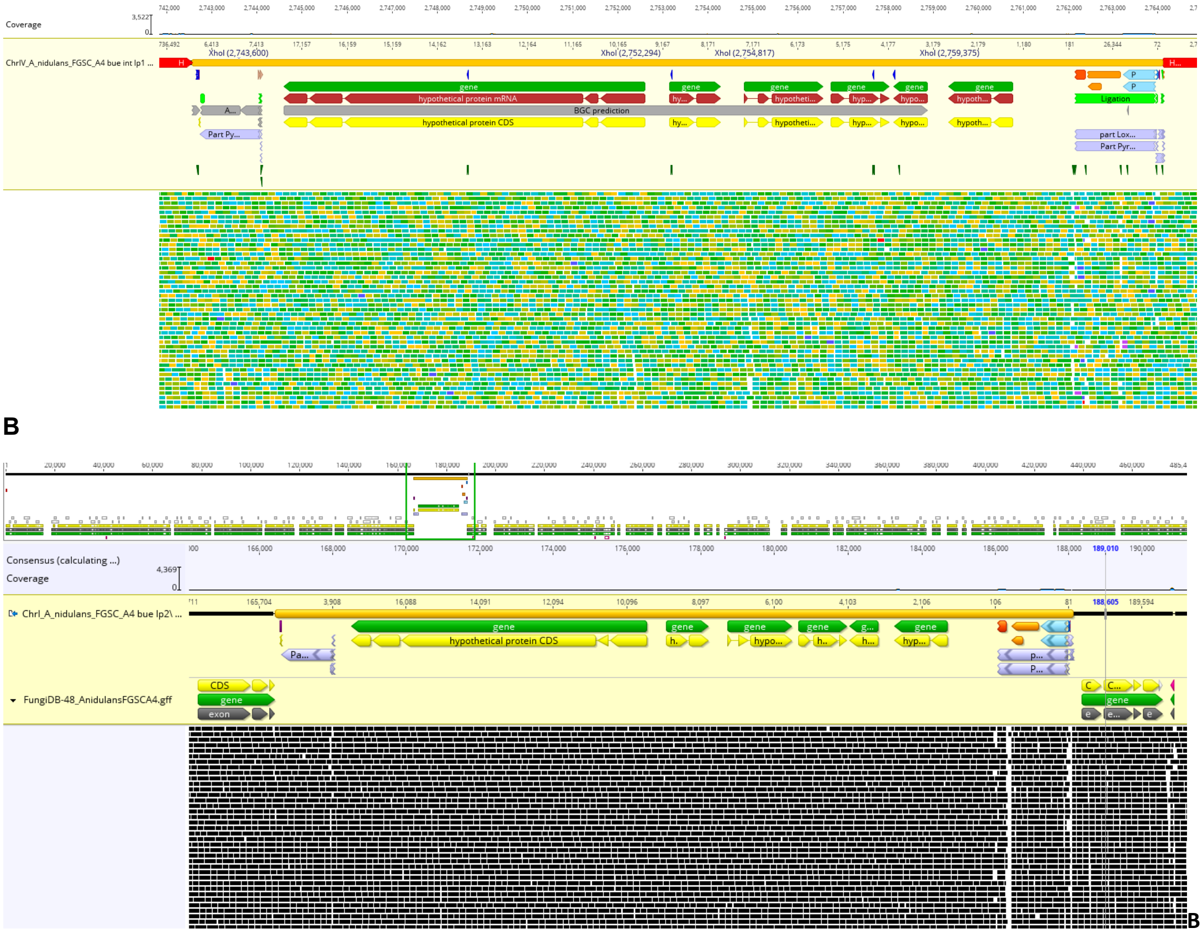


**Figure S10.** Chromosomal integration of *bue* genes at LP1 (**A**) and LP2 (**B**) observed by whole genome sequencing. Biological duplicate from with *bue* genes integrated at LP2 also demonstrate de expected recombination event and no mutations on *bue* genes that could explain the lack of compound production from a chromosomal context. Low quality alignments on the right region of the figure are caused by the presence of *A. nidulans* $P_{gpdA}$ and $T_{trpC}$ in *mCherry* expression cassette. Figure visualized in Geneious 11.03.

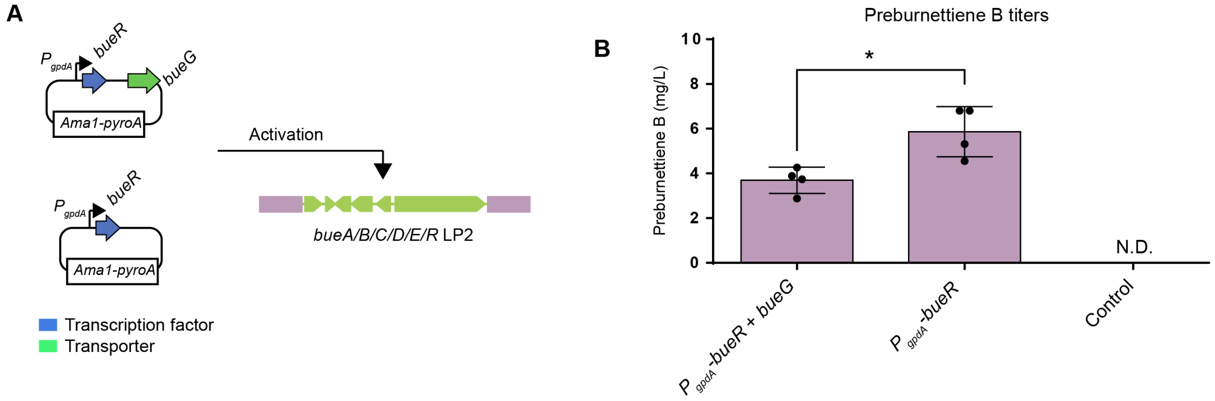


**Figure S11.** The transcription factor BueR is responsible for the activation of the *bue* biosynthetic gene cluster. **A.** Schematic of the vectors evaluated containing the transcription factor encoding gene *bueR* under the promoter *gpdA* ($P_{gpdA}$) alone or with the presence of the transporter encoding gene *bueG*. **B.** Preburnettiene B (**1**) titers from liquid culture extracts from transformants grown in GMM supplemented with uracil, uridine and riboflavin (only selective for *pyroA* marker). Interestingly, there is a significative increase in the production of **1** in the absence of *bueG*. Values are the mean of four biological replicates, specific values are indicated as black dots, and error bars represent standard deviation. Two-sided Welch's T-test p-value = 0.014. N.D.= non-detectable.

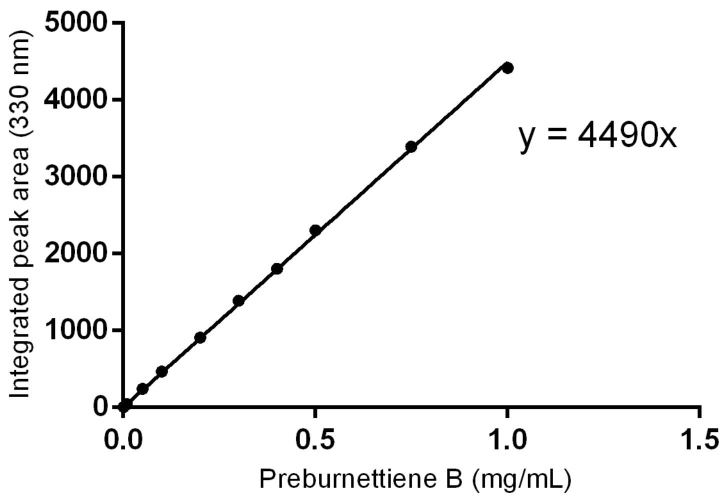


**Figure S12.** Calibration curve used to quantify the production of preburnettiene B (**1**). The regression coefficient was used to estimate the concentration in the crude extract and then extrapolate to the titer of **1** per liter of culture media.

### Supporting Tables

**Table S1** DNA sequence of *lox* recombination sites used, 5′ to 3′. The nucleotides that differ from the original *loxP* sequence are underlined. The symbol used for each *lox* site is indicated at the right column.

| Site | Left end | Core | Right end | Symbol |
|---|---|---|---|---|
| *loxP* | ATAACTTCGTATA | ATGTATGC | TATACGAAGTTAT | 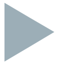 |
| *lox2272-71* | ATAACTTCGTATA | AAGTATCC | TATACGAACGGTA | 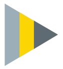 |
| *lox2272-66* | TACCGTTCGTATA | AAGTATCC | TATACGAAGTTAT | 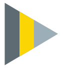 |
| *lox2272-72* | TACCGTTCGTATA | AAGTATCC | TATACGAACGGTA | 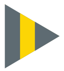 |
| *lox2272* | ATAACTTCGTATA | AAGTATCC | TATACGAAGTTAT | 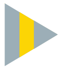 |

**Table S2** Vectors used in this work.

| Vector name | Purpose and vector short name in the text | Origin | Vector Size (kb) | Addgene ID |
|---|---|---|---|---|
| pGem-PgpdA-Cre-TtrpC | Helper vector for the expression of Cre. Not replicative on *A. nidulans*. | This work | 4.9 | 168784 |
| pGemLP1-loxP-bar-Lox2272-71 | Vector to create LP1 by homologous recombination. | This work | 6.6 | 168785 |
| pGemLP2-loxP-bar-Lox2272-71 | Vector to create LP2 by homologous recombination. | This work | 6.6 | 168786 |
| pGem-loxp-mcherry-lox66-2272 | Donor_vector-1 | This work | 6.6 | - |
| pRecomb-loxp-mcherry-4Gcloningsite-lox66-2272 | Donor_vector-2 | This work | 12.2 | - |
| pRecomb-loxp-mcherry-bueABCDER-lox66-2272 | Donor_vector-2-*bue* | This work | 27.3 | 168787 |
| pKW20088-loxP-mcherry-pyrG-lox66-227266 | Comparison to mCherry episomal expression | This work | 16.2 | - |
| pYFAC-bueABCDER | AMA1-based expression of bueA/B/C/D/E/R (*pyrG* marker). | [7] | 32.3 | - |
| pYFAC-bueG-PgpdA-bueR | Transcription factor overexpression for *bue* cluster activation (*pyroA* marker) and transporter expression. | [7] | 19.9 | - |
| pYFAC- PgpdA-bueR | Transcription factor overexpression for *bue* cluster activation (*pyroA* marker). | This work | 16.9 | - |

**Table S3** Genotype of parental *Aspergillus nidulans* strains used in this work.

| Parental strains | Relevant genotype |
| --- | --- |
| *A. nidulans* LO8030 [8] | pyroA4, riboB2, pyrG89, nkuA::argB, Sterigmatocystin cluster (*AN7804–AN7825*)Δ, emericellamide cluster (*AN2545-AN2549*)Δ, asperfuranone cluster (*AN1039–AN1029*)Δ, monodictyphenone cluster (*AN10023–AN10021*)Δ, terraquinone cluster (*AN8512–AN8520*)Δ, austinol cluster part1 (*AN8379–AN8384*)Δ, austinol cluster part2 (*AN9246–AN9259*)Δ, f9775 (*AN7906–AN7915*)Δ, asperthecin cluster (*AN6000–AN60002*)Δ |
| LO8030-LP1 | LO8030 + loxP-bar-lox2272-71::Sterigmatocystin cluster (*AN7804–AN7825*)Δ |
| LO8030-LP2 | LO8030 + loxP-bar-lox2272-71::IS1 |
| LO8030-LP1-bueA/B/C/D/E/R | LO8030 + loxP-mCherry-bueA/B/C/D/E/R-pyrG-lox2272-72::Sterigmatocystin cluster (*AN7804–AN7825*)Δ |
| LO8030-LP2-bueA/B/C/D/E/R | LO8030 + loxP-mCherry-bueA/B/C/D/E/R-pyrG -lox2272-72::IS1 |

**Table S4.** Oligonucleotides and gBlock used in this work.

| Oligonucleotide | Sequence | Purpose |
|---|---|---|
| Cre-Lic-FW | TAGAGGATCACTCGTCGTGTCGCCCATGTCTAATTTGTTGA CTGTTCATCAAAATT | Cloning Cre into pBargpe1 |
| Cre-Lic-Rv | GCCCCCCCTCGAACGTCGTCGTCGCCCATCACCATCTTCC AAGAGTCTAACC | Cloning Cre into pBargpe1 |
| RvSp_i147 | ACGTCGTCGTCGCCCTAATCACCATCTTCCAAGAGTCTAAC C | To build pGem-PgpdA-Cre |
| FwSp_147 | UAGGGCGACGACGACGTTCG | To build pGem-PgpdA-Cre |
| i147_AW_Fw | ACCGTCCGTCTCTCCGCATGTTCACTGGCCGTCGTTTTACA AC | To build pGem-PgpdA-Cre |
| i147_AW_Rv | ACAGCTCATCTGCAATGCATCATGGTCATAGCTGTTTCCTG TG | To build pGem-PgpdA-Cre |
| TtrcRv | ATGCATTGCAGATGAGCTGTATC | To build pGem-PgpdA-Cre |
| PgpdA-Fw | CATGCGGAGAGACGGACGG | To build pGem-PgpdA-Cre |
| AattI-Lox2272-71Rv | ATTATAGATCATAACTTCGTATAAAGTATCCTATACGAACGG TAGCTAGCTGTACGT | To build pGemLP1-loxP-bar-Lox2272-71 |
| AattI-Lox2272-71Fw | ACAGCTAGCTACCGTTCGTATAGGATACTTTATACGAAGTT ATGATCTATAAT | To build pGemLP1-loxP-bar-Lox2272-71 |
| FwUpST | TTCCTCGTCGTGTCGCCCAAGTGTAGGTAATGATTAGGATT GCTGG | To build pGemLP1-loxP-bar-Lox2272-71 |
| RvUpST | CCGCTCATGAGACAATAACCCTGTCCCACCTTGATCTACCT GC | To build pGemLP1-loxP-bar-Lox2272-71 |

| | | |
|---|---|---|
| FwDownST | TATACATCGCAGGGGGTTGACCAAAGGTTGGGTCGTCGGTA | To build pGemLP1-loxP-bar-Lox2272-71 |
| RvDownST | CGTCGTCGTCGCCCAGCATTGGACAGCGCGTTAAT | To build pGemLP1-loxP-bar-Lox2272-71 |
| FwpBarLoxPLox2272 | AGGGTTATTGTCTCATGAGCGG | To build pGemLP1-loxP-bar-Lox2272-71 |
| RvpBarLoxPLox2272 | GTCAACCCCCTGCGATGTATA | To build pGemLP1-loxP-bar-Lox2272-71 |
| i150_HA2TS1_Fw | ATATTAAGGGTTCCGGATCGCGGCGGTTGATTGAGTCGAGATAGTC | To build pGemLP2-loxP-bar-Lox2272-71 |
| i150_HA2TS1_Rv | CATATGGTCGACCTGCAGGCGGCCGCCCAAGTTGTCGGAAGGATGACC | To build pGemLP2-loxP-bar-Lox2272-71 |
| i150_HA1TS1_Rv | TCCGCTCATGAGACAATAACCCTTTACACAGACCAGCAATCATCAAAC | To build pGemLP2-loxP-bar-Lox2272-71 |
| i150_HA1TS1_Fw | CATGCTCCCGGCCGCCATGGCGGCCGCGGACATCTCTCATGGCTCGC | To build pGemLP2-loxP-bar-Lox2272-71 |
| gBlock 1 | CATCGCCGGTCGAGGCATCTACGCTGCTCCCGACCCGGTTGAAGCTGCACAGCGGTACCAGAAAGAAGGCTGGGAAGCTTATATGGCACGTGTCTGTGGTAAAAGCTGAACTCTGATTGCTAGCCAGAACTACCGTTCGTATAAAGTATCCTATACGAAGTTATCCCTTAAGGTATATGCGGCAAGTCATGATTTCCTCTTGGAGCAAAAGTGTAGTGCCAGTACGAGTGTTGTGGAGGAAGGCTGCATACATTGTGCCT | To build pKW20088-loxP-mcherry-pyrG-lox2272-66 |

| | | |
|---|---|---|
| LoxP-pyrG-F | TAACTTCTTCGGCGACAGCATCACCGACTTCGTTAATTAATGGGCTCTGAGTGTGTTTGG | To build pKW20088-loxP-mcherry-pyrG-lox2272-66 and amplifying fragments for *in vitro* recombination |
| pyrG-LoxP-R | GTCTTTGGCAACACCAAACACACTCAGAGCCCATTAATTAACGAAGTCGGTGATGCTGTC | To build pKW20088-loxP-mcherry-pyrG-lox2272-66 |
| pKW-LoxP-F | AGTAACCTCGCGGGTGTTCTTGACGATGGCATCCTGCGACGTCGACTCTAGAGGATCCCC | To build pKW20088-loxP-mcherry-pyrG-lox2272-66 |
| PyrgPartR | GCCATATAAGCTTCCCAGCCT | To build pKW20088-loxP-mcherry-pyrG-lox2272-66 |
| LIC-mcherry-FW | CTCGTCGTGTCGCCCATGGTGAGCAAGGGCGAGGAG | To build PgpdA-mCherry-TrpcT |
| LIC-mcherry-Rv | CGTCGTCGTCGCCCATTACTTGTACAGCTCGTCCATGC | To build PgpdA-mCherry-TrpcT |
| pacIgpdeRv | ATGTCTTTGGCAACACCAAACACACTCAGAGCCCATTAATTAACGTCATGCATTGCAGAT | To build pGem-loxp-mcherry-lox71-2272 and pKW20088-loxP-mcherry-pyrG-lox2272-66 |
| pacIgpdeFw | TCTTAACTTCTTCGGCGACAGCATCACCGACTTCGCGATCGCATGCGGAGAGACGGACG | To build pGem-loxp-mcherry-lox71-2272 and pKW20088-loxP-mcherry-pyrG-lox2272-66 |
| NotI-PkwF | TACATCATGCGGCCGCCACTGAGAACCATGGCACCGAAG | To build pGem-loxp-mcherry-lox71-2272 |
| NotI-Gblock-Rv | TACTATAAGCGGCCGCCTTACCTCACTTGCTAGATGACTGG | To build pGem-loxp-mcherry-lox71-2272 |
| PyrgTBsmbIRv | CAATCACGCATTCCCGAGTG | To build pRecomb-loxp-mcherry-4Gcloningsite-lox71-2272-66 |
| pyrgT-mut-bsmbI-F | AATGGATACGTGCACTCGGGAATGCGTGATTGGCTCGACTCGTGTTCTCAGTTCCTCATTCC | To build pRecomb-loxp-mcherry-4Gcloningsite-lox71-2272-66 |
| Crelox-4g-Fw | ATCTGCAATGTTAAGAATTCGCTGATTGTGATAGTTCCCACTTG | To build pRecomb-loxp-mcherry-4Gcloningsite-lox71-2272-66 |

| | | |
|---|---|---|
| Crelox-4g-Rv | CACACTCAGAGCCCAGAATTCTGATCTTCTCATCACGCCTC TTG | To build pRecomb-loxp-mcherry-4Gcloningsite-lox71-2272-66 |
| Crelox-mche-Fw | AGATGCGTAAGGAGAGTCGACTCTAGAGGATCCCCG | To build pRecomb-loxp-mcherry-4Gcloningsite-lox71-2272-66 |
| Crelox-mche-Rv | CACAATCAGCGAATTCTTAACATTGCAGATGAGCTGTATCT GG | To build pRecomb-loxp-mcherry-4Gcloningsite-lox71-2272-66 |
| Crelox-ori-Fw | AGAAGATCAGAATTCTGGGCTCTGAGTGTGTTTGGTG | To build pRecomb-loxp-mcherry-4Gcloningsite-lox71-2272-66 |
| Crelox-ori-Rv | TCCTCTAGAGTCGACTCTCCTTACGCATCTGTGCG | To build pRecomb-loxp-mcherry-4Gcloningsite-lox71-2272-66 |
| Bur_140_Fw1 | CCTCTTCCAGATACAGCTCATCTGCAATGTTAACGTGCGAC CCTGAAACTACA | To build pRecomb-loxp-mcherry-BueABCDER-lox71-2272-66 |
| Bur_Rv1 | AACCGTTGGGCGACAAGG | To build pRecomb-loxp-mcherry-BueABCDER-lox71-2272-66 |
| Bur_Fw2 | AGGCTCATCTTCCTTGTCGC | To build pRecomb-loxp-mcherry-BueABCDER-lox71-2272-66 |
| Bur_Rv2 | GTCGCTCTCGATGATGGTACAC | To build pRecomb-loxp-mcherry-BueABCDER-lox71-2272-66 |
| Bur_Fw3 | CGTAGGCCGAGTGTACCATC | To build pRecomb-loxp-mcherry-BueABCDER-lox71-2272-66 |
| Bur_Rv3 | GGGTTGCCTGTCTGTCCC | To build pRecomb-loxp-mcherry-BueABCDER-lox71-2272-66 |

| | | |
|---|---|---|
| Bur_Fw4 | GCGGACGTTATCTTTGGGGAC | To build pRecomb-loxp-mcherry-BueABCDER-lox71-2272-66 |
| Bur_140Rv | TTTGGCAACACCAAACACACTCAGAGCCCAATGCTTGCGAATGGTGCACC | To build pRecomb-loxp-mcherry-BueABCDER-lox71-2272-66 |
| BueR_Rv | GATGAGACCCAACAACCATGATACCAGGGGGCAAGTTGTACCCCTTTGAACAGCTGGTTACTCTGTTGGTTTCTCCACGG | To build pYFAC- PgpdA-bueR |
| HA1TS1_Scr | GGCTTGGACATCAAATGGGC | Screening PCR |
| HA2TS1_Scr | AACACCCTGGATTTCGTGGG | Screening PCR |
| BarDownstream-F-Seq | GACGTGGGTTTCTGGCAGC | Screening PCR 1 |
| PbarUpstream-R-Seq | GCATTCATTGTTGACCTCCACTAGC | Screening PCR 2 |
| Gblock-R | CTTACCTCACTTGCTAGATGACTGG | Amplifying fragments for *in vitro* recombination |
| P1FwUpST | GACGACGAATACTATGCATCCTTG | Screening PCR 2 and 4 (LP1) |
| P6RVDownST | CAGAGGACCCCAGGAACGA | Screening PCR 1 and 3 (LP1) |
| RMCEgpda_Fw | CCGGGTACTCGCTCTACCTAC | Screening PCR 3 |
| RMCEpyrg_RV | TCCAAAGGATCGCTGGCTAC | Screening PCR 4 |
